# (-)-Englerin A binds a conserved lipid site of TRPC5 and exposes a Met-aromatic motif in channel activation

**DOI:** 10.1101/2025.07.09.663840

**Authors:** Sebastian A. Porav, Alexandra Ptakova, Claudia C. Bauer, Kasia L. R. Hammond, David J. Beech, Viktorie Vlachova, Stephen P. Muench, Robin S. Bon

## Abstract

TRPC4/5 cation channels are polymodal cellular sensors that play key roles in signal transduction/integration and have been implicated in various human pathologies, including anxiety, pain and cardiometabolic disease^1–3^. The plant natural product (-)-englerin A (EA)^4^ is a potent, selective TRPC4/5 agonist^5,6^ that has transformed fundamental and translational research on TRPC4/5 channels. However, the structural basis of interactions between EA and TRPC4/5 proteins has remained elusive, limiting our ability to fully understand and exploit mechanisms of TRPC4/5 channel activation by this intriguing natural product. Here, we present nine high-resolution cryo-EM structures (2.4-3.2 Å) of human TRPC5 – representing different states and ligand occupancies – which show that EA binds to a conserved lipid binding site between transmembrane domains of adjacent TRPC5 subunits. Our structural models are consistent with the effects of mutagenesis of nearby residues on EA’s potency, efficacy and activation kinetics, and allow us to rationalise competitive inhibition by other TRPC4/5 modulators as well as EA’s selectivity profile within the TRPC family. Comparison of structures containing various TRPC5:EA stoichiometries revealed key structural and molecular determinants of EA-mediated TRPC5 activation – most notably the aromatic interaction network around Phe520 – underscoring the critical function of Met-aromatic motifs in ion channel structure and function. Binding of EA causes conformational changes of nearby amino acid residues, resulting in rearrangement of the pore helices into a pre-open state. Collectively, we provide structural insight into the mode-of-action of the most widely used TRPC4/5 agonist, which will underpin fundamental TRPC4/5 channel research as well as ongoing drug discovery programmes.

## Main

The 28 mammalian Transient Receptor Potential (TRP) proteins form tetrameric, non-selective cation channels^7–9^. These TRP channels are exquisite cellular sensors that mediate cellular responses to external stimuli such as temperature, pH, force, metal ions, lipids, and small molecules^10^. Many TRP channels are modulated by plant natural products present in food and herbal medicines^11,12^, for example capsaicin (red hot chili pepper; TRPV1^13,14^), cannabidiol (cannabis; TRPV2^15,16^), piperlongumine (long pepper; TRPV2^17^), menthol (mint; TRPM8^18–20^), allyl isothiocyanate (mustard; TRPA1^21,22^), cinnamaldehyde (cinnamon; TRPA1^21^) and galangin (galangal; TRPC5). Conversely, such phytochemicals have critically enabled the study of structure, function and biological relevance of specific TRP channels^11,23–25^.

The guaiane sesquiterpene derivative (-)-englerin A (EA) was first isolated from the stem bark of *Phyllanthus engleri*^26^, which has been used as both traditional medicine and poison in southern Africa^4^. The discovery that EA selectively kills A498 renal cancer cells^26^ sparked major research into the elucidation of its chemical structure, absolute configuration and molecular target(s), and the synthesis and biological characterisation of EA and its derivatives^4,27^. Mechanism-of-action studies revealed that EA is a potent (nanomolar) and selective TRPC4/5 channel agonist, and that TRPC4/5 channels are essential mediators of the cytotoxic effect of EA on A498 and other TRPC4/5-expressing cancer cells, and of its adverse effects upon administration in mice^5,6,28,29^. These discoveries resulted in the widespread use of EA as a chemical probe of TRPC4/5 channels in biological studies and drug discovery^2,30^.

TRPC4 and TRPC5, which readily form homo-and heteromeric cation channels (especially with the widely expressed modulatory subunit TRPC1) are key mediators of cellular responses in the brain/central nervous system, gut and cardiovascular system^1,2,30^. The implication of TRPC4/5 channels in various human diseases and disorders, including seizures, fear-related behaviour, pain, heart failure and cardiometabolic disease^1–3,30^, has led to drug discovery programmes targeting TRPC4/5 as well as clinical trials with TRPC4/5 inhibitors for the treatment of anxiety/post-traumatic stress disorder (BI 1358894)^31^, kidney disease (GFB-887)^32^ and peripheral neuropathy (ONO-2910).

Cryogenic electron microscopy (cryo-EM) has enabled the determination of high-resolution structures of TRPC4^33–37^ and TRPC5^36,38–43^ channels, providing key insights into channel architecture and interaction with TRPC1, metal ions, lipids, other proteins, and drug-like small-molecule modulators. However, despite major efforts, the molecular mechanism by which EA selectively activates TRPC4/5 channels has remained elusive. To address this knowledge gap, we used cryo-EM to determine high-resolution cryo-EM structures of the human TRPC5 channel (2.4-3.2 Å) in complex with EA. Our structures show that EA binds to a conserved lipid binding site, consistent with its pharmacological profile and the effect of mutagenesis of nearby residues. By comparing structures containing various TRPC5:EA stoichiometries and supported by functional characterisation of TRPC5 variants through intracellular calcium recordings and patch-clamp electrophysiology, we provide structural insight into molecular determinants of EA-mediated TRPC5 activation. Structural similarity between TRPC1, TRPC4 and TRPC5, and the conservation of the lipid/EA/xanthine binding site, suggest that our results extrapolate more generally to the entire TRPC1/4/5 sub-family, while comparison to TRPC3/6/7 allows us to rationalise EA’s selectivity profile. Our detailed understanding of the interaction of this therapeutically relevant class of ion channels with its most widely used agonist will be critical for ongoing TRPC4/5 drug discovery programmes.

### EA binds to a conserved lipid binding site of TRPC5

Several observations suggest that EA occupies a well-defined ligand binding site of TRPC4/5 channels: 1) excised membrane patch recordings in the presence or absence of G protein blockade suggest direct and reversible TRPC4/5 channel activation by EA via a site accessible via the external membrane leaflet^5^; 2) mutagenesis results in TRPC5 variants that respond to EA with altered potency and biophysical characteristics^39,44,45^; and 3) the EA response of TRPC4/5 channels is competitively inhibited by other small molecules, such as the xanthine Pico145 and the EA analogue A54^46–48^. However, attempts by us and others^40^ to obtain cryo-EM structures of TRPC5:EA complexes by incubation of purified TRPC5 protein with excess EA resulted in empty ‘apo’ structures. We hypothesised that this outcome reflected the limited solubility of EA. Addition of the carrier pluronic acid (PA) to EA preparations allows consistent channel activation in cellular studies^5,46^. Therefore, we decided to obtain the structure of TRPC5 in the presence of both EA and PA (‘TRPC5:EA^PA^’). To ascribe possible structural modifications induced by PA, we also obtained a new ‘apo’ TRPC5 structure in the presence of only PA (‘TRPC5^PA^’). We used our C-terminally truncated, MBP-tagged human TRPC5 construct (MBP-PreS-hTRPC5_Δ766-975_), which is biochemically stable, retains its pharmacological response to EA, and has been used for the determination of multiple high-resolution TRPC5 structures^36,39,43^.

We determined two distinct structures of TRPC5:EA^PA^, representing different channel states within the same sample, using 3D variability analysis (3DVA)^49^ in CryoSparc^50^ (**Supporting Note 1**). The structures of these states, TRPC5:EA^PA^-S1 and TRPC5:EA^PA^-S2, were both resolved to 2.5 Å (C4 symmetry) (**Figure 1a-f; Extended Data Figure 1; Extended Data Figure 2**). TRPC5:EA^PA^-S1 and TRPC5:EA^PA^-S2 mainly differ in terms of the rotational position of their intracellular ankyrin repeat domains (ARDs) and coiled-coil domains (CCDs) (**Extended Data Figure 1d-g; Supporting Note 1**), and resemble previously reported TRPC5 structural states (**Supporting Note 1**)^40,42^. In addition, we determined the structure of ‘apo’ TRPC5^PA^ to a resolution of 2.4 Å (C4 symmetry) (**Figure 1g-i; Extended Data Figure 1; Extended Data Figure 2**), representing ARD state 2. To exclude heterogeneity between TRPC5 subunits, all maps were also refined without imposed symmetry (C1), resulting in near-identical structures, albeit with lower global resolutions by ∼0.3 Å. The high-resolution, high-quality C4 maps allowed us to build structural models de novo using ModelAngelo^82^, followed by manual curation and refinement of the models (**Figure 1; Extended Data Figure 3**).

**Figure 1.**
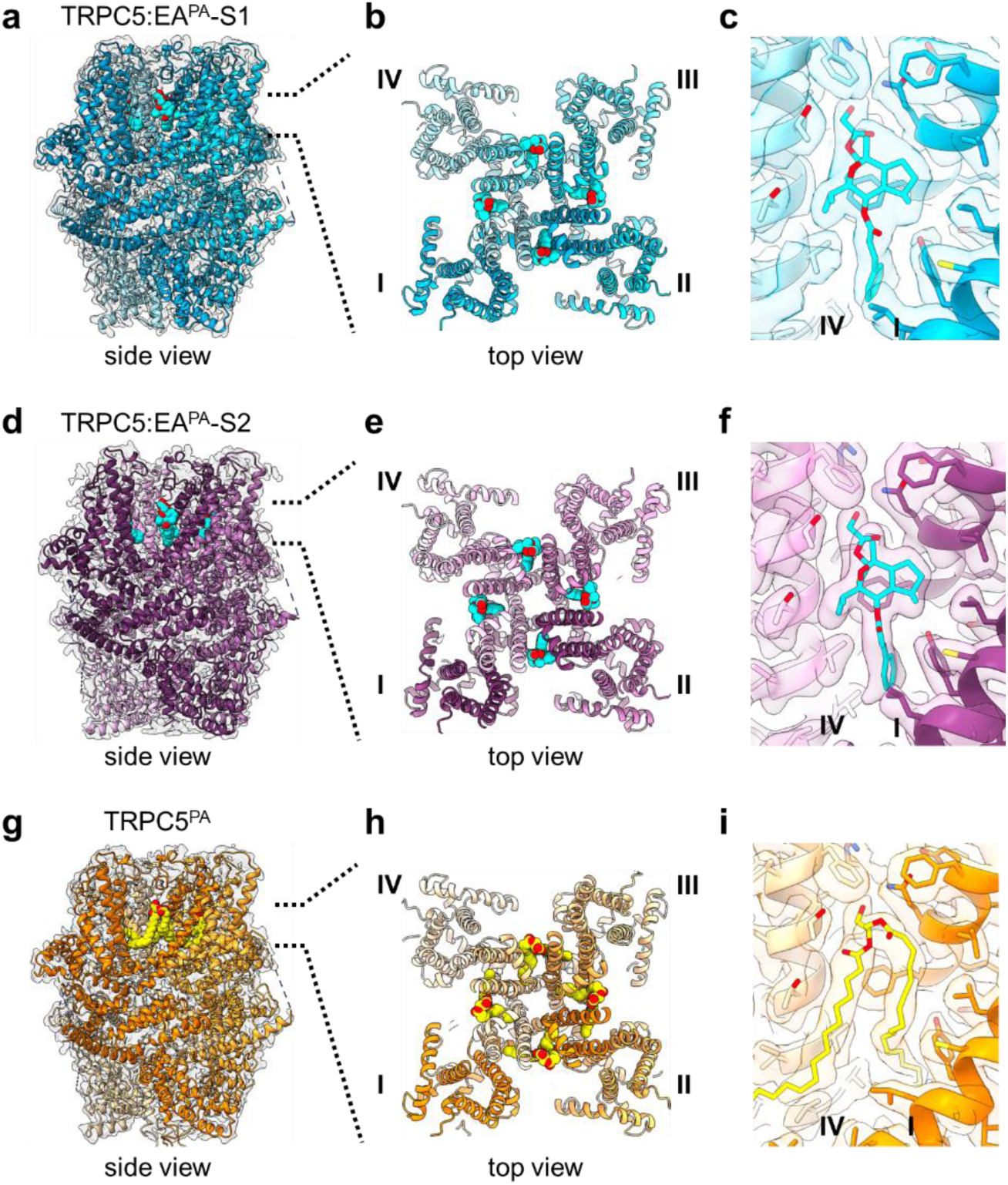
Cryo-EM structures reveal the EA binding site of the human TRPC5 channel. a,b, 3D cryo-EM map and model of TRPC5:EA^PA^-S1 as seen from the side (a) and top (b). The four modelled TRPC5 subunits (I-IV) are shown as ribbons in shades of blue in the grey EM map; EA is shown in cyan (carbon atoms) and red (oxygen atoms). **c**, Close-up of the EA binding site between TRPC5 subunits I and IV in (a,b). **d,e,** 3D cryo-EM map and model of TRPC5:EA^PA^-S2 as seen from the side (d) and top (e). The four modelled TRPC5 subunits (I-IV) are shown as ribbons in shades of magenta in the grey EM map; EA is shown in cyan (carbon atoms) and red (oxygen atoms). **f**, Close-up of the EA binding site between TRPC5 subunits I and IV in (d,e). **g,h,** 3D cryo-EM map and model of TRPC5^PA^ as seen from the side (g) and top (h). The four modelled TRPC5 subunits (I-IV) are shown as ribbons in shades of orange in the grey EM map; the resident lipid (modelled as 1-oleoyl-2-palmitoyl-*sn*-glycerol consistent with recently published TRPC5 structures^40,42^) is shown in yellow (carbon atoms) and red (oxygen atoms). **i**, Close-up of the lipid binding site between TRPC5 subunits I and IV in (g,h). In all three maps, we found a few small, disordered regions in the cytosolic domains, for which poor density impaired model building; the corresponding residues are listed in **Extended Data Table 2**.

### General Architecture

Our TRPC5 structures are of high quality (**Extended Data Table 1; Extended Data Figure 2; Extended Data Figure 3**) and their overall architectures resemble previously reported ones^36,38–43^, with dimensions of 100 Å × 100 Å × 120 Å and characterised by a two-layer architecture: the intracellular cytosolic region (ICR) and the transmembrane region (TMD) (**Figure 1a,d,g**). The intracellular region is formed by the N-terminal ankyrin repeat domain (ARD), a linker-helix domain (LHD), pre-S1 elbow, TRP domain, connecting helix, and CCD of each monomer (**Extended Data Figure 1, Figure 3c**). The transmembrane domain comprises six transmembrane helices (**Extended Data Figure 3**), the first four of which (S1-S4) assemble into a voltage sensor-like domain (VSLD). The last two transmembrane helices (S5 and S6) and the re-entrant pore helix (E3) from each monomer establish the channel pore. Comparison of the TRPC5^PA^ map to those of previously determined ‘apo’ TRPC5 maps^38–40^ confirmed that that addition of PA did not affect the overall TRPC5 structure.

### The EA binding site

Comparison of the TRPC5:EA^PA^-S1 and TRPC5:EA^PA^-S2 maps to the TRPC5^PA^ map (and to previously determined ‘apo’ TRPC5 maps^38–40^) revealed clear differences in the non-protein densities in the conserved lipid binding sites^39^ located between adjacent TRPC5 monomers. The TRPC5^PA^ map displays a well-defined, characteristic lipid U-shape (**Figure 1i**), whereas the corresponding sites in the TRPC5:EA^PA^-S1 and TRPC5:EA^PA^-S2 maps contain densities closely resembling the shape and size of EA (**Figure 1; Extended Data Figure 1h**). C1 reconstructions revealed consistent density across all four sites, consistent with full occupancy of four molecules of EA per TRPC5 tetramer. Therefore, we focused our further analysis on the C4 reconstructions. The quality of our maps and the high local resolution (2.2-2.4 Å) allowed us to unambiguously model EA into each of the four ligand binding sites (**Figure 1; Extended Data Figure 1h**). For clarity, we labelled the four TRPC5 monomers (I-IV) in a counterclockwise manner (seen from the extracellular side; **Figure 1**), consistent with Won et al.^37^ **Figure 1c,f,i** show the EA/lipid binding site between TRPC5 monomers I and IV, which is formed by S5 and the pore re-entrant helix of TRPC5(I) and S6 of TRPC5(IV). The binding site is highly hydrophobic, with two small hydrophilic regions around EA’s 7-isopropyl group and its five-membered ring (**Extended Data Figure 1i**).

### Molecular interactions between EA and TRPC5

Modelled binding interactions between TRPC5 and EA are near-identical in TRPC5:EA^PA^-S1 and TRPC5:EA^PA^-S2 (**Figure 2a,b**). EA interacts with TRPC5 protein through a hydrogen bond network involving the hydroxyl group of EA’s glycolate substituent, the carboxamide of Gln573(I)’s side chain and – in TRPC5:EA^PA^-S1 – the indole NH of Trp577(I). In addition, the phenyl ring of EA’s 6-cinnamate substituent forms a π–π stacking interaction with the side chain of Tyr524(I). Analysis of the hydrophobic environment shows contacts between the 6-cinnamate and nearby residues from both TRPC5 monomers, specifically Val610(IV), Val614(IV), Phe520(I), Tyr524(I) and – in TRPC5:EA^PA^-S2 – Leu521(I). The hydrophobic landscape is further enhanced by contacts between Phe576(I) and EA’s 7-membered ring, as well as between Leu528(I) and EA’s 5-membered ring. The intricate network of hydrophobic contacts stabilising the TRPC5:EA complex partially mimics the interactions of the resident lipid in TRPC5:EA^PA^ (**Figure 2c**) and in previously reported TRPC5 ‘apo’ structures^38–40^.

**Figure 2.**
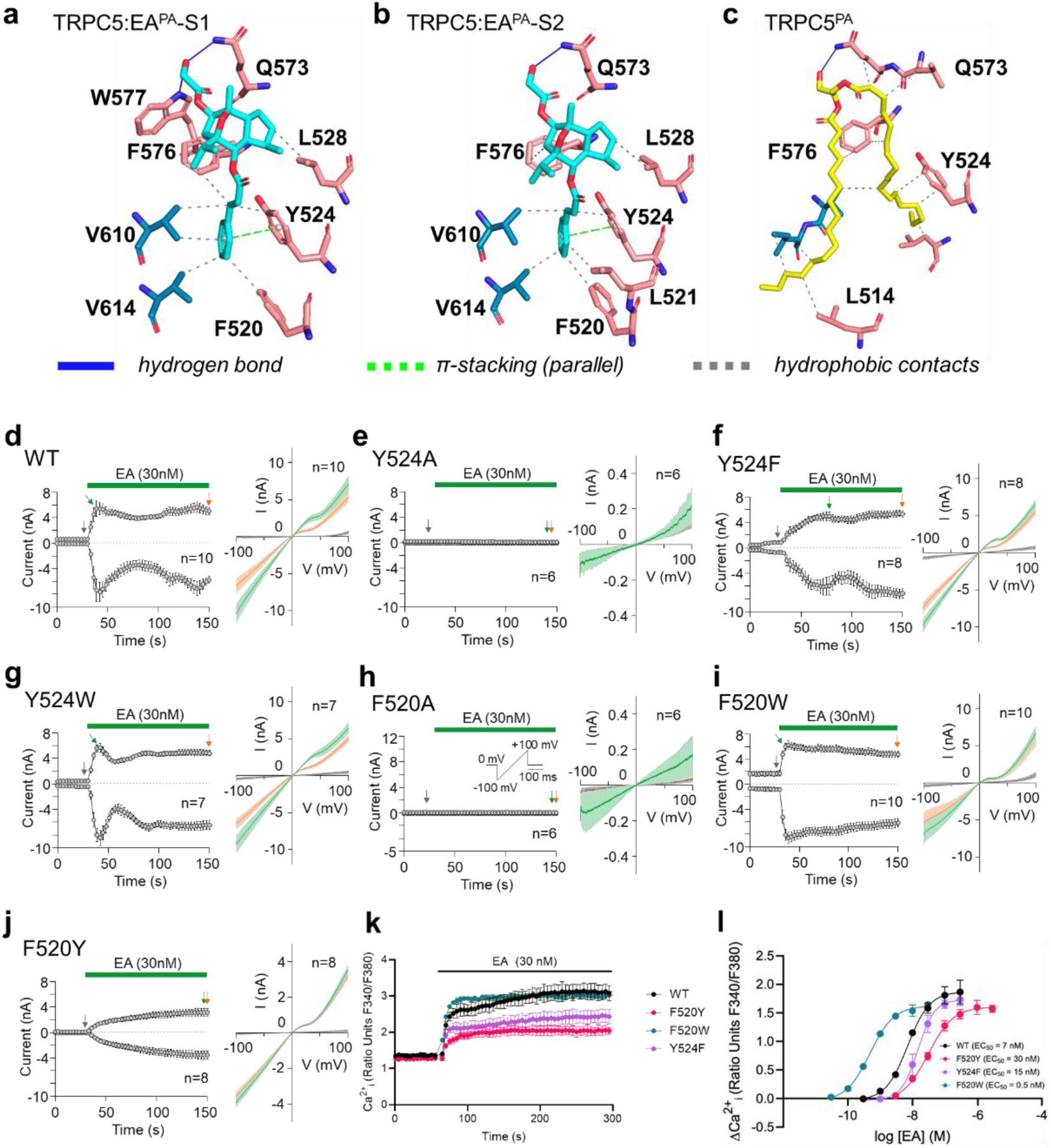
The TRPC5:EA binding model is consistent with effects of site-directed mutagenesis of binding site residues. a-b. Key TRPC5 residues involved in interactions with EA (cyan) in TRPC5:EA^PA^-S1 (a) and TRPC5:EA^PA^-S2 (b). **c,** Key TRPC5 residues involved in interactions with the resident lipid (yellow) in TRPC5^PA^. Hydrogen bonds (blue), π-stacking interactions (green) and hydrophobic contacts (grey) are annotated in panels a-c. Residues from monomer I are shown in pink and residues of monomer IV are shown in blue. **d-j,** Time courses of average whole-cell currents elicited by 30 nM EA in HEK293T cells expressing indicated TRPC5 constructs. A ramp pulse from-100 mV to +100 mV was periodically applied from a holding potential of 0 mV every 3 seconds for 500 ms (shown in panel h). Amplitudes were measured at-100 mV and +100 mV and the mean ± SEM was plotted as a function of time. On the right-hand side of each panel, mean current-voltage relations (coloured curves, ± SEM as lighter-coloured envelopes) are plotted for the currents measured at times indicated on the left-hand side of the panel by vertical arrows (grey at baseline, green at peak, and orange after 2-min exposure to EA). The number of cells (n) is indicated. Summary data from patch-clamp experiments are plotted as bar graphs in **Extended Data Figure 5a-c**. **k,** Representative recordings from one 96-well plate (N=6) showing Ca^2+^ influx in response to 30 nM EA in HEK293 cells expressing TRPC5-SYFP2 variants. Data shown as mean ± SD **l,** Concentration-response data from intracellular Ca^2+^ recordings for TRPC5-SYFP2 mutants transiently expressed in HEK293 cells. Cells were stimulated with EA (0.03 nM to 3 µM) to activate TRPC5 channels and Ca^2+^ influx was measured to determine EC_50_ values. Data shown as mean ± SEM (n/N=3/18). EC_50_ values were determined with a four-parameter non-linear regression fit. Representative data traces for EC_50_ determinations are shown in **Extended Data Figure 6g-j**.

**Figure 3.**
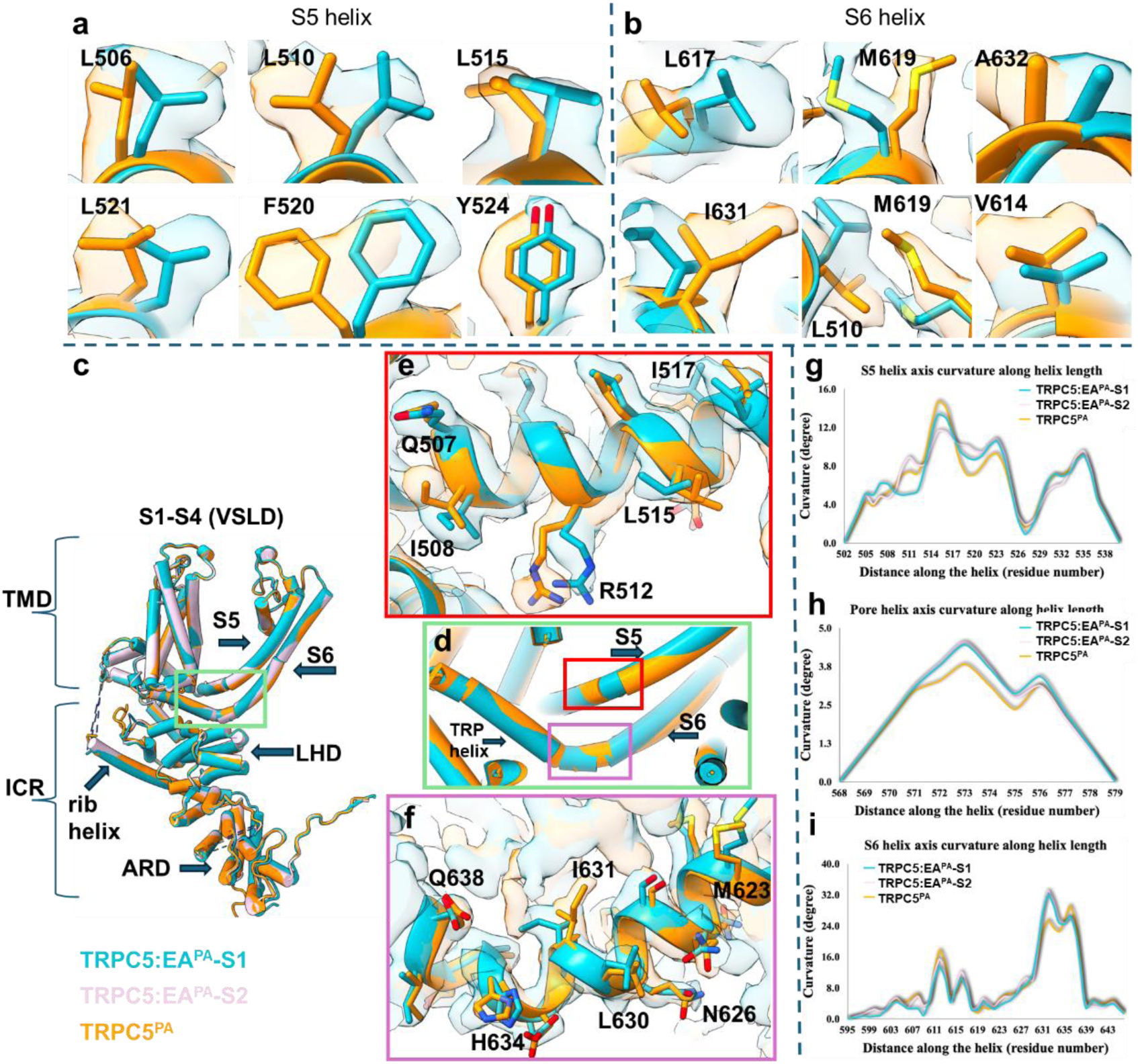
EA binding induces conformational changes of TRPC5 residues and helices. a-b, Close-ups of superimposed maps and models TRPC5:EA^PA^-S1 (cyan) and TRPC5^PA^ (orange) showing conformational changes of TRPC5 residues on the S5 helix and S6 helix, respectively. **c,** Superimposed model of monomers of TRPC5:EA^PA^-S1 (cyan), TRPC5:EA^PA^-S2 (pink), and TRPC5^PA^ (orange), highlighting the transmembrane domain (TMD), intracellular cytosolic region (ICR), voltage sensor-like domain (VSLD) composed of transmembrane helices S1-S4, rib helix, linker helix domain (LHD), ankyrin repeat domain (ARD) and the pore helices (S5 and S6). **d-f,** close-up of domains (maps and models) of TRPC5:EA^PA^-S1 (cyan) and TRPC5^PA^ (orange) showing the changes in curvature of the S5 and S6 helices. **g-i,** Curvature analysis (using HELANAL-Plus software^54^) of TRPC5 helices involved in the formation of the EA binding site: the S5 helix (g), the pore re-entrant helix (h), and the S6 helix (i).

Our EA binding model allows us to rationalise the effects of mutagenesis of specific TRPC5 residues on EA-mediated channel activation. We previously showed that mutation of Gln573, Phe576 or Trp577 results in a substantial drop in EA potency (i.e. 46-fold for TRPC5_Q573T_, >246-fold for TRPC5_F576A_ and 65-fold for TRPC5_W577A_)^39^. Our structures show that these residues make direct interactions with EA.

In addition, replacement of Gly606 – located one helix turn upstream of Val610 – by tryptophan completely prevented channel activation by 30 nM EA without compromising activation by positive voltage^45^. Here, we further tested the functional effects of mutagenesis of key EA interacting residues of TRPC5. For functional analysis of TRPC5 variants, we tested EA responses in patch-clamp recordings from HEK293T cells (**Figure 2d-j**; **Figure 5n; Extended Data Figure 4** and quantified data from all independent experiments in **Extended Data Figure 5a-c**). We also tested responses of all TRPC5 variants to voltage activation (**Figure 4d-f**; **Figure 5l,m; Extended Data Figure 4a**). For selected TRPC5 variants, we performed surface biotinylation assays (to test whether non-functional channels can still insert into the plasma membrane; **Extended Data Figure 5d**) and intracellular calcium recordings with various agonists (**Figure 2k,l; Extended Data Figure 6**).

**Figure 4.**
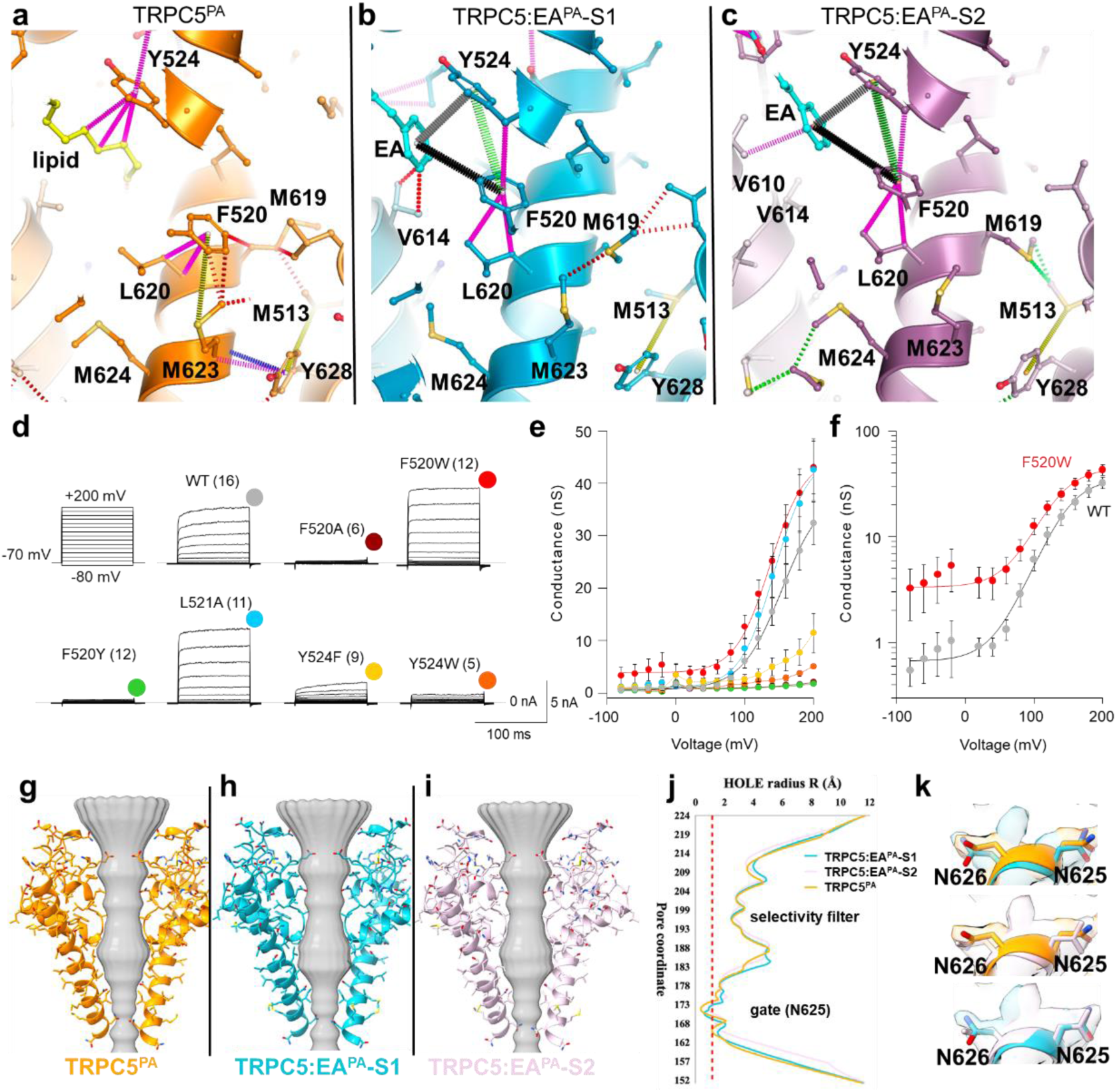
Changes in the aromatic landscape around Phe520 mediate TRPC5 channel gating. a-c, Aromatic interactions around residue Phe520 in TRPC5^PA^ (orange), TRPC5:EA^PA^-S1 (cyan) and TRPC5:EA^PA^-S2 (pink). Methionine sulfur-π interactions are shown in yellow, carbon – π interactions in magenta, and different geometries of ring interactions are shown in green (OF), black (ET) and grey (FT), as defined in Jubb et al.^59^ **d,** Mean current traces in response to 100-ms voltage steps from-80 to +200 mV (protocol shown top left) recorded from HEK293T cells expressing indicated TRPC5 constructs. The currents were recorded ∼1 min after whole-cell patch formation in extracellular control solution. Numbers of cells (n) are indicated in parentheses. **e,** Average conductances obtained from steady-state currents measured at the end of the pulses as indicated by coloured symbols atop the records in (d). Solid lines in wild-type TRPC5 (WT), TRPC5_F520W_ and TRPC5_L521A_ are best fits to a Boltzmann function as described in the Materials and Methods. In less responsive mutants, the lines connecting the data points have no theoretical meaning. **f**, A semi-logarithmic plot of the average steady-state conductance-voltage relationships measured from wild-type TRPC5 and TRPC5_F520W_ as in panel (d). At negative potentials, TRPC5_F520W_ exhibits a disturbed closed–open equilibrium in favour of the open state. **g-j,** Pore shape analysis for TRPC5:EA^PA^-S1 (light blue), TRPC5:EA^PA^-S2 (pink) and TRPC5^PA^ (orange) calculated using PoreAnalyzer^60^; in (j), the pore radius is plotted against the pore coordinate down the channels, highlighting the selectivity filter and lower gate. The red dashed line indicates where the pore becomes too narrow for water to pass through (1.2 Å), showing that the three structures represent closed states. **k,** Close-up of gatekeeper residue Asn625 with overlayed EM map (mesh) and model of TRPC5:EA^PA^-S1 (light blue), TRPC5:EA^PA^-S2 (pink) and TRPC5^PA^ (orange) showing minor changes and the presence of additional density in EA maps suggesting a degree of flexibility.

**Figure 5.**
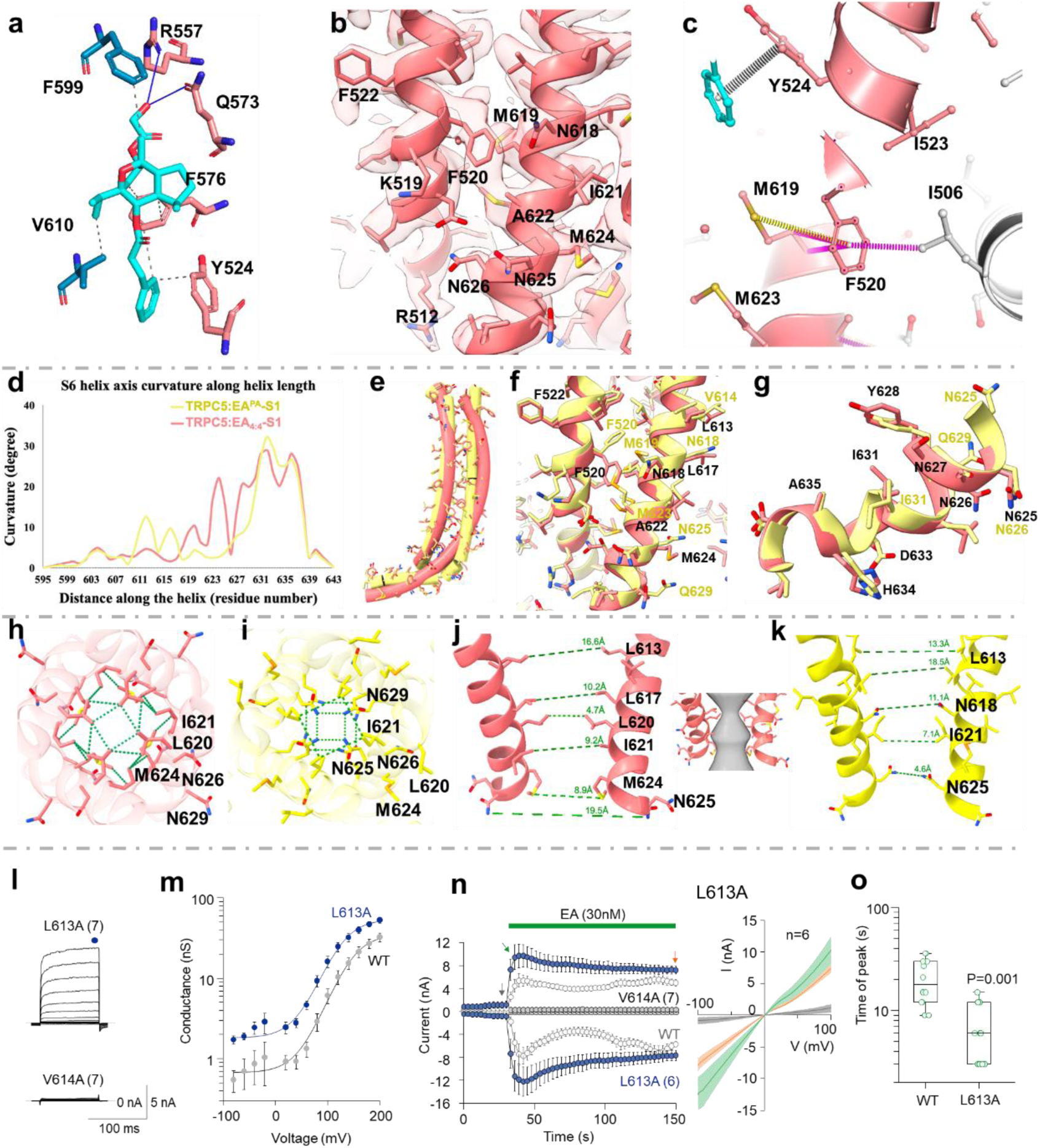
Transition of the TRPC5 π-bulge to an α-helix is implicated in EA-mediated channel gating. a, Interactions of EA with TRPC5 amino acid residues in TRPC5:EA**_4:4_**-S1 (blue line indicated hydrogen bonds while the grey dotted line shows hydrophobic contacts). **b,** The domain surrounding Phe520 in TRPC5:EA**_4:4_**-S1 (model as ribbons in the EM map), showing a new conformation of Phe520 compared to TRPC5 structures described above. **c,** Aromatic interactions around residue Phe520 in TRPC5:EA**_4:4_**-S1. TRPC5 subunits are shown in pink (subunit I) and grey (subunit IV). The phenyl ring of EA’s 6-cinnamate substituent is shown in cyan. Methionine sulfur-π interactions are shown in yellow, carbon – π interactions in magenta, and different geometries of ring interactions are shown in grey (FT), as defined in Jubb et al.^59^ **d,e,** Comparative analysis of S6 helix curvature in TRPC5:EA**_4:4_**-S1 (red) and TRPC5:EA^PA^-S1 (yellow). Curvature was calculated using HELENAL-Plus. **f,g,** Close-ups of the S5 and S6 helices in (e) showing the rearrangements of Phe520 and surrounding residues (f) and the π-bulge to α-helix transition (g). **h,i,** Formation of a hydrophobic gate in TRPC5:EA**_4:4_**-S1 (h) compared with the ionic gate found in TRPC5:EA^PA^-S1 (i). **j,k,** Pore rearrangement and formation of a hourglass-shaped gate in TRPC5:EA**_4:4_**-S1 (j) as a result of a register shift downwards from L613 as compared to TRPC5:EA^PA^-S1 (k). **l,** Average current traces in response to a voltage step protocol (from-80 to +200 mV; 20 mV increments) recorded from HEK293T cells expressing TRPC5_L613A_ or TRPC5_V614A_. **m,** The average steady-state conductance-voltage relationship for TRPC5_L613A_ and wild-type TRPC5 (WT), measured as in panel (l). **n,** Time-course of average whole-cell currents elicited by 30 nM EA in HEK293T cells expressing indicated TRPC5 constructs. A ramp pulse from-100 mV to +100 mV was periodically applied from a holding potential of 0 mV every 3 seconds for 500 ms. Left, currents were measured at-100 mV and +100 mV and the mean ± SEM was plotted as a function of time. Numbers of cells (n) are indicated. Right, mean current-voltage relations (coloured curves, ± SEM as lighter-coloured envelopes) plotted for the TRPC5_L613A_ currents measured at times indicated in the left panel by vertical arrows (grey at baseline, green at peak, and orange after 2-min exposure to EA). Number of cells (n) is indicated. **o,** Box plot of mean values of time to maximal current mediated by wild-type TRPC5 and TRPC5_L613A_ within 2 min of 30 nM EA application, measured at +100 mV. The box denotes the 50^th^ percentile (median) as well as the 25^th^ and 75^th^ percentile. The whiskers mark the 5^th^ and 95^th^ percentiles. The indicated probability was obtained from the Student’s two-sided unpaired t-test that was performed in order to determine if there was a significant difference between the values for the two variants. Summary data from patch-clamp experiments are plotted as bar graphs in **Extended Data Figure 5a-c**.

We first focused on the two aromatic residues that interact with EA. Mutation of Tyr524 to alanine rendered the channels (TRPC5_Y524A_) non-functional; the variant did not respond to EA, voltage activation or any other tested activators (**Figure 2e; Extended Data Figure 6**), even though cell surface expression of the channel protein was not affected (**Extended Data Figure 5d**). Compared to wild-type TRPC5 (**Figure 2d**), variant TRPC5_Y524L_ produced small and slowly developing currents in response to 30 nM EA (**Extended Data Figure 4b**) and a minimal voltage response (cf. **Figure 4d** and **Extended Data Figure 4a**). Substitution of Tyr524 with an aromatic residue phenylalanine (TRPC5_Y524F_) retained the maximum response to EA, albeit with significantly slower activation kinetics (**Figure 2f; Extended Data Figure 5b**). In contrast, replacement of Tyr524 with the larger and more hydrophobic aromatic residue tryptophan (TRPC5_Y524W_) produced functional channels with a significantly faster onset of EA activation compared to wild-type TRPC5 (**Figure 2g; Extended Data Figure 5b**). Variant TRPC5_F520A_ was unresponsive to EA (**Figure 2h**), voltage stimulation (**Figure 4d**) or other tested agonists (**Extended Data Figure 6**) without affecting surface expression (**Extended Data Figure 5d**). Currents through TRPC5_F520W_ showed significantly faster responses to 30 nM EA compared to wild-type channels (**Figure 2i, Extended Data Figure 5b**), whereas TRPC5_F520Y_ showed significantly slower activation and exhibited lower amplitudes of responses within 2 minutes of EA application (**Figure 2j, Extended Data Figure 5b,c**). Although TRPC5_F520Y_ could be activated by EA, it lost its response to AM237^51^, BTD^52^, high extracellular Ca^2+^,^53^ or carbachol in combination with Gd^3+^ (**Extended Data Figure 6**). Variant TRPC5_F520L_ produced nonfunctional channels (**Extended Data Figure 4a,c; Extended Data Figure 6**). These data are consistent with the importance of Y524 and F520 in EA-mediated TRPC5 activation. This was further confirmed by intracellular calcium recordings, in which responses of TRPC5 variants to 30 nM EA matched those obtained in patch-clamp experiments (**Figure 2k**). Additional concentration-response measurements showed that mutation of Y524 or F520 to other aromatic residues does gives similar maximum efficacy, but results in EC_50_ shifts by up to an order of magnitude (**Figure 2l; Extended Data Figure 6g-i**).

We then examined the requirement of hydrophobic residues at positions 521, 610 and 614 for channel activation by EA. Variant TRPC5_L521A_ resulted in significantly slower activation kinetics of the EA-evoked recurrent (**Extended Data Figure 4d, Extended Data Figure 5b**). However, within 2 min of EA application, this response still reached a mean amplitude that was not statistically significantly different from wild-type channels at +100 mV (**Extended Data Figure 5c**), indicating that EA efficacy was preserved in this channel variant. This result suggests that leucine’s bulky, hydrophobic side chain is important for the interaction between TRPC5 and EA. TRPC5_L521F_ was constitutively active at negative potentials (**Extended Data Figure 4e**), with EA activation kinetics similar to wild-type. This is possibly due to phenylalanine’s larger, aromatic side chain, which could introduce new stabilising interactions or alter conformational equilibria, favouring an active-like state even in the absence of an agonist. Variants TRPC5_V610A_ (**Extended Data Figure 4f**) and TRPC5_V614A_ (**Figure 5n**) showed no or very small responses to EA (< 22.4 and 24.8 pA/pF, respectively), but the conservative mutation TRPC5_V610L_ produced robust EA-induced currents (**Extended Data Figure 4g**).

Overall, these functional data are consistent with the EA binding site and binding mode in our TRPC5 structures (**Figure 1**; **Figure 2**) and with the physico-chemical properties of the molecular interactions between EA and TRPC5.

### EA induces conformational changes of key TRPC5 domains

#### The pore helices

Because the residues forming the EA binding site originate from three different TRPC5 helices – S5, S6, and the re-entrant pore helix – we examined their architecture in greater detail. In TRPC5:EA^PA^-S1 and TRPC5:EA^PA^-S2, the structures of these domains are near-identical, but distinct from those in the ‘apo’ structure TRPC5:EA^PA^ (**Figure 3**). Upon EA binding, the side chains of Leu617 and Met619 move away from and towards the symmetry axis, respectively (**Figure 3b**). The most notable change is in the rotameric switch of Phe520, which moves 2.2 Å (measured from the ipso carbon of the phenyl ring) from an orientation that is transversal and away from the symmetry axis to one that points upwards and towards the symmetry axis (**Figure 3a**). This rearrangement likely favours interaction of Phe520 with EA’s 6-cinnamate substituent (**Figure 2a,b**). Although Phe520 and Met619 are located on different helices, their side chains are proximal and face each other (**Figure 4a-c**). The movements of these two side chains appear to occur synergistically and in a co-dependent manner to initiate a chain reaction that propagates downwards through both S5 and S6, causing rearrangements of adjacent residues and ultimately affecting the helix curvature (**Figure 3c-i**).

#### The aromatic interaction network

Closer analysis of Phe520 revealed that EA induces changes in its aromatic interaction network^55,56^ (**Figure 4**). In the ‘apo’ structure TRPC5:EA^PA^, Phe520 engages in two types of π interactions: a C/π interaction with Leu620 and a more notable S/π interaction with Met623 (**Figure 4a**). Near Phe520, we also find other ring systems, specifically Tyr524 from the same chain and Tyr628 from the adjacent chain. Examination of the connections between these systems revealed that Tyr524 interacts with the lipid tail through a C/π interaction. Similarly, Tyr628 forms a C/π interaction with Met623, which is involved in the S/π interaction with Phe520 and engages in an S/π interaction with Met513. Binding of EA disrupts the S/π interactions between Phe520 and Met623, leading to the incorporation of Phe520 into a new three-ring interaction system (**Figure 4b,c**). This new geometry involves Tyr524, Phe520, and EA’s 6-cinnamate substituent. The repositioning of the Phe520 ring facilitates the formation of a C/π interaction with Leu620 as well as with Tyr524, while changes induced by EA binding do not affect the S/π system between Met513 and Tyr628.

Methionine S/π interactions are known for stabilising protein structures^57^, but their functional roles are poorly understood. Therefore, we explored the functional role of this aromatic interaction network in detail using mutagenesis. As described above, the aromaticity of Phe520 is essential for TRPC5 activation by EA, voltage or other agonists (**Figure 2h-l**; **Figure 4d-f; Extended Data Figure 4a**). While TRPC5_F520A_ is non-functional, EA can activate TRPC5_F520Y_ and TRPC5_F520W_, albeit with altered potency and activation kinetics (**Figure 2d,h-l; Extended Data Figure 5b**). An impaired kinetic fingerprint is also seen with variants TRPC5_L620A_ (**Extended Data Figure 4a,h**) and TRPC5_L620F_ (**Extended Data Figure 4a,i**), supporting the importance of the Phe520-Leu620 interaction. The TRPC5 channel is considered intrinsically voltage-sensitive because it can be activated by positive voltages (> + 60 mV) in the absence of any agonists^58^. Variant TRPC5_F520W_ exhibited a gain-of-function phenotype (**Figure 4d-f**), suggesting that tryptophan at this position reduces the activation energy for channel opening. Upon voltage stimulation, this construct exhibited tonic activation at negative membrane potentials and a leftward shift of the half-maximal activation voltage (*V*_50_) by about 15 mV (139.6 ± 5.3 mV versus 155.9 ± 4.3 mV for wild-type; *P* = 0.024; *n* = 12 and 16). The apparent number of gating charges, *z*, was 0.87 ± 0.04 *e*_o_, which is not different from wild-type channels (0.84 ± 0.03 *e*_o_; *P* = 0.531), indicating that the mutation TRPC5_F520W_ lowers the activation threshold for channel activation, but it does not affect the voltage-sensing mechanism itself.

Phe520 is fully conserved in human TRPC channels, and highly conserved throughout the entire human TRP family (**Extended Data Figure 5e**), suggesting an important role in TRP(C) channel function. Likewise, Met623 is fully conserved in the TRPC family while Met619 is replaced by a leucine in TRPC1 (**Extended Data Figure 5e**). Alanine substitutions at these two methionine residues (TRPC5_M619A_ and TRPC5_M623A_) strongly suppressed EA-and voltage-induced currents and the deleterious effect was even more pronounced in TRPC5_M623L_ (**Extended Data Figure 4a,j-l**). In intracellular calcium recordings, a response of TRPC5_M619A_ could be seen when a much higher EA concentration (1 µM) was applied (**Extended Data Figure 6b**). Among the amino acids analysed, Tyr524 is the least conserved in the TRPC family, being found only in TRPC4/5, and replaced by phenylalanine in other TRPC members (**Extended Data Figure 5e**). Substitution of Tyr524 by phenylalanine (TRPC5_Y524F_) or tryptophan (TRPC5_Y524W_) led to dramatically decreased currents evoked by depolarising voltage (**Figure 4d,e**). Surprisingly, the activation kinetics of EA responses in TRPC5_Y524W_ were significantly faster than those of wild-type channels (**Figure 2d,g; Extended Data Figure 5b**), suggesting that Tyr524 plays a dual role in TRPC5 gating by participating in voltage-and EA-dependent activation pathways that do not fully overlap. In contrast, substitution of the nearby residue Leu521 by alanine (TRPC5_L521A_) completely preserved voltage activation (**Figure 4d,e**) but drastically slowed down EA activation (**Extended Data Figure 4d; Extended Data Figure 5b**). These selective effects of mutations on individual modalities of TRPC5 activation are indicative of a separate and likely conserved mechanism of voltage-dependent gating. Overall, these insights suggest that EA-or voltage-induced changes to the aromatic interaction network around Phe520 and Tyr524 are critical to TRPC5 channel activation.

#### The channel pore

EA is a high-efficacy agonist of TRPC5, and our structures show full occupancy of TRPC5 as well as conformational changes upon EA binding. But how does EA binding affect the structure of the TRPC5 pore? Compared to the ‘apo’ structure TRPC5^PA^, the pore radius doubles in TRPC5:EA^PA^-S1 and nearly triples in TRPC5:EA^PA^-S2 (**Figure 4g-j**). However, this expansion is still insufficient for the passage of water molecules or ions. Closer inspection of the local maps surrounding the gatekeeper residue Asn625 revealed an additional density between Asn625 and Asn626, which could not be modelled (**Figure 4k**). This density may represent water molecules in a standby mode (i.e. waiting for ions to pass through the channel) that are involved in the rewatering process. Further investigation of the maps along the ion conduction path revealed several lipid-like densities (or structured water molecules) and two bulkier densities located in the selectivity filter and hydrophobic gate regions, which – given the composition of buffers used – are most likely Na^+^ ions. These additional densities were consistently observed in all three maps, so their presence is unrelated to EA binding. Furthermore, the high resolution of the obtained maps allowed us to model several water molecules both around and within the protein. These results suggest that our EA-bound TRPC5 structures represent non-conducting states, albeit with clear changes to pore residues.

#### The S6 π-bulge

In various TRP channels, the π-bulge within the S6 helix has been implicated in channel modulation^61^. As EA binds proximal to the S6 π-bulge, we analysed this region in greater detail. In the map of ‘apo’ TRPC5^PA^, the entire S6 helix is well-resolved, allowing unambiguous modelling of the π-bulge around Leu613 and Val614 (**Extended Data Figure 7**). In contrast, the maps obtained in the presence of EA (TRPC5:EA^PA^-S1 and TRPC5:EA^PA^-S2) displayed relatively low local resolution in this area, resulting in imperfect fits of Leu613 and Val614 (**Extended Data Figure 7**). In the presence of EA, the S6 helix could be manually modelled in two different states, containing either a π-bulge or an extended α-helix. Notably, this conformational heterogeneity of S6 has not been observed in any of the TRPC1/4/5 structures published so far, suggesting that this region is important for EA-induced channel gating.

### Structures with various TRPC5:EA stoichiometries reveal additional channel states

#### TRPC5:EA structures in the absence of PA

To further test the effects of the addition of PA to our samples, including on local maps and on channel states, we decided to try an alternative sample preparation method. Instead of adding an EA/PA preparation to TRPC5 after purification, we maintained EA (100 µM) in all solutions throughout the sample preparation process, from cell lysis to grid making. Because the resulting cryo-EM map suggested partial occupancy of TRPC5 by EA, we used 3DVA and clustering to separate our data set, allowing us to determine six distinct TRPC5:EA structures, which we categorised based on the ARD rotational state and the TRPC5:EA binding stoichiometry (**Supporting Note 2; Extended Data Figure 8; Extended Data Figure 9**). Models were built from all six maps, and quality of the models is illustrated in **Extended Data Figure 10** and **Extended Data Table 1**. Further analysis of the effects of TRPC5:EA binding stoichiometry focused on two of the ‘ARD state 1’ TRPC5 structures: TRPC5:EA**_4:4_**-S1 (with four EA molecules bound per TRPC5 tetramer) and TRPC5:EA**_4:2_**-S1 (with two EA molecules bound per TRPC5 tetramer).

#### Rearrangement of the S5 and S6 pore helices

Because both TRPC5:EA^PA^-S1 and TRPC5:EA**_4:4_**-S1 contain TRPC5:EA at 4:4 stoichiometry, we initially attempted to fit the TRPC5:EA^PA^-S1 model into the map of TRPC5:EA**_4:4_**-S1. Visual inspection of the fitting accuracy revealed several inconsistencies, particularly around the S6 π-bulge, the pore gate and the EA binding site. In addition, the EA binding site contained some lipid density, which remained present in all four binding sites upon map refinement without applying symmetry (C1) or with C4 relaxed symmetry. This suggests the inclusion of a small number of particles with incomplete EA occupancy in our final map, which we do not consider to significantly impact the overall structure and model. To create an accurate model of TRPC5:EA**_4:4_**-S1, we manually rebuilt the regions that did not fit well. Comparison of this model to that of TRPC5:EA^PA^-S1 revealed a global RMSD of 1.6 Å, indicating that TRPC5:EA**_4:4_**-S1 represents a new conformation of the TRPC5:EA complex.

Inspection of the EA binding sites of TRPC5:EA**_4:4_**-S1 resulted in the identification of two additional residues that can interact with EA through hydrophobic contacts and hydrogen bonding: Phe599(I) and Arg557(IV), respectively (**Figure 5a**). Notably, Phe520 did not interact with EA in this TRPC5 channel conformation (**Figure 5a-c**). In TRPC5:EA**_4:4_**-S1the benzyl group of Phe520 is rotated by ∼125° compared to TRPC5:EA^PA^-S1, pointing downward and away from the symmetry axis of the channel (**Figure 5a-c**). This position of Phe520 was not seen in any of the previously described structures. Additional weak density around Phe520 suggests some degree of flexibility (**Figure 5b**). When exploring the aromatic interactions of Phe520, we found that its conformation is stabilised by S-π and C-π interactions with Met619 from the same subunit, as well as C-π interactions with Ile506 from the adjacent subunit (**Figure 5c**).

Further comparison of TRPC5:EA**_4:4_**-S1 to TRPC5:EA^PA^-S1 revealed clear differences in helix curvature, with a local RMSD of the S6 helices of 2.9 Å. This is the result of a full transition of the S6 π-bulge into an α-helical structure, as determined by the rotameric change of Leu613 and rearrangement of Val614 (**Figure 5d-f**). This transition induces clockwise rotation of the S6 helix, resulting in the shifting of each of residues Leu613-Tyr628 by one position. Consistent with these observations, we found that TRPC5_V614A_ resulted in no or very small responses to voltage (< 5.4 nS at +200 mV), EA (< 24.8 pA/pF) (**Figure 5l,n**) or other activators (**Extended Data Figure 6c-f**), although the protein expressed at the plasma membrane (**Extended Data Figure 5d**) to form channels that gave responses when a higher concentration of EA was applied (**Extended Data Figure 6b**). In contrast, TRPC5_L613A_ showed a gain-of-function phenotype (**Figure 5l-o**). This construct was constitutively active at negative potentials and – upon voltage stimulation – displayed a leftward shift of *V*_50_ to 135.7 ± 6.2 mV (*P* = 0.014; *n* = 7) (**Figure 5l-m**). The apparent number of gating charges, *z*, was not different from wild-type channels (0.88 ± 0.04 *e*_o_; *P* = 0.492), which means that the mutation does not affect the intrinsic voltage-sensing mechanism of the channel, but makes it easier to activate by shifting the activation voltage to more negative values. Accordingly, the onset kinetics of EA responses mediated through TRPC5_L613A_ were significantly faster compared with wild-type channels at both negative and positive potentials (median of the time of peak outward current 6 s; 3–15 s; versus 21 s; 9–36 s for wild-type TRPC5) (**Figure 5n,o; Extended Data Figure 5b**).

The S6 rearrangement of TRPC5:EA**_4:4_**-S1 also determines its shortening by three residues, extending the connecting loop between S6 and the TRP-helix from Asp633-Ala635 in TRPC5^PA^ and TRPC5:EA^PA^-S1 to Leu630-Ala635 in TRPC5:EA**_4:4_**-S1 (**Figure 5g**). Likewise, EA binding not only affects the curvature of the S6 helix but, as the main constituent of the pore, changes the physicochemical properties of the pore through its rearrangement. The S6 rotation starting from Leu613 induces an upward shift of the gate position and a change of gate type from an ionic gate to a hydrophobic gate (**Figure 5h,i**). This hydrophobic gate formation at Leu620 is promoted by the entrance of the Leu620 side chain into the pore – pushing Ile621 to the side – and the replacement of Asn625, in the centre of the pore, by Met624 (**Figure 5h-k**). Pushing outward of the Asn625 residues enlarges the pore diameter, distal to Leu620, resulting in the formation of a pseudo-symmetrical hourglass shape (**Figure 5j,k**). The bottleneck found at Leu620 suggests that this residue may act as a lever in the opening and closing of the channel. Analysis of hydrophobic contacts around the new channel gate illustrates an intricate mesh-like interaction network between Leu620, Ile621, and Met624 (**Figure 5h,i**). We hypothesise that the conformational state described here represents a desensitised state or a transitional state after ion passing.

### Partial EA binding results in asymmetric, intermediate channel states

We obtained two maps (one for each ARD state) that represent TRPC5 tetramers with two EA molecules bound in opposing binding sites, and lipids in the other two opposing binding sites (TRPC5:EA**_4:2_**-S1 and TRPC5:EA**_4:2_**-S1; **Figure 6; Extended Data Figure 8; Extended Data Figure 9**). These structures uniquely allow comparison of the effects of binding of EA vs lipid within the same TRPC5 tetramer. We observed no major differences between the two maps apart from the rotation of the ARD domain, so we focus our analysis on TRPC5:EA**_4:2_**-S1. This structure contains remarkable features in terms of the EA/lipid binding sites (including Phe520’s aromatic interaction network), the S5 and S6 helices, and the channel pore (**Figure 6**; **Figure 7**).

**Figure 6.**
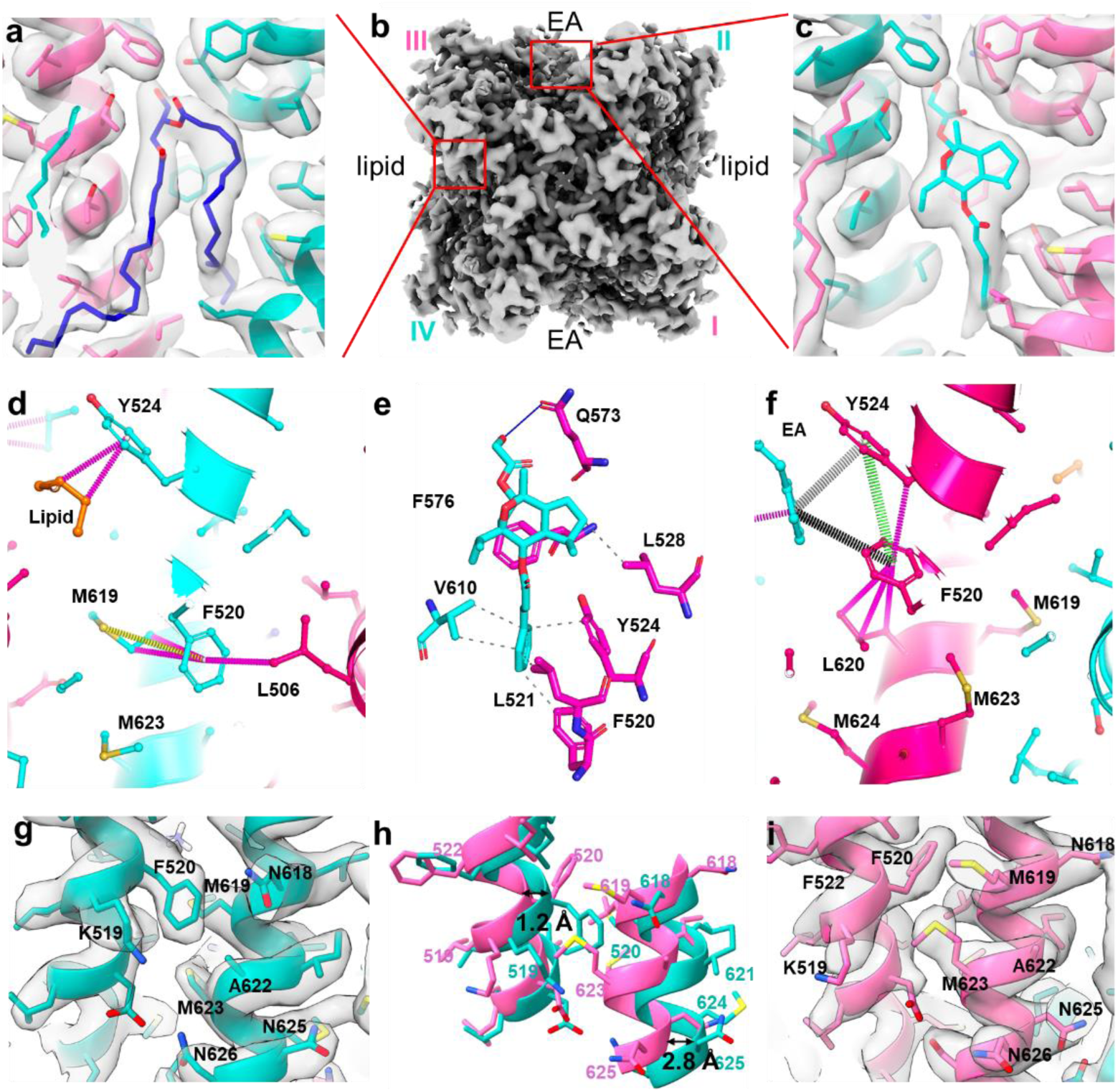
Partial EA binding results in an asymmetric, intermediate TRPC5 channel state. a-c, cryo-EM map and model of TRPC5:EA_4:2_-S1 (b) showing the alternating occupancy of TRPC5 binding sites by lipid (blue) in (a) and EA (cyan) in (c). TRPC5 subunits I and III are displayed in pink and TRPC5 subunits II and IV in cyan. **d-f,** Analysis of the different lipid/EA binding sites in TRPC5:EA**_4:2_**-S1. Key interactions of EA (cyan) with residues from TRPC5 subunits I (pink) and IV (cyan) are displayed in (e). Major differences in the aromatic interactions networks of Phe520 are visible between sites occupied by lipid (orange) in (d) and those occupied by EA (cyan) in (f). **g-i,** partial cryo-EM map and model of S5 and S6 helices (around Phe520) from TRPC5 subunits IV (cyan; g) and I (pink; i) and model superimposition of the two subunits (h) showing rearrangements caused by EA binding.

**Figure 7.**
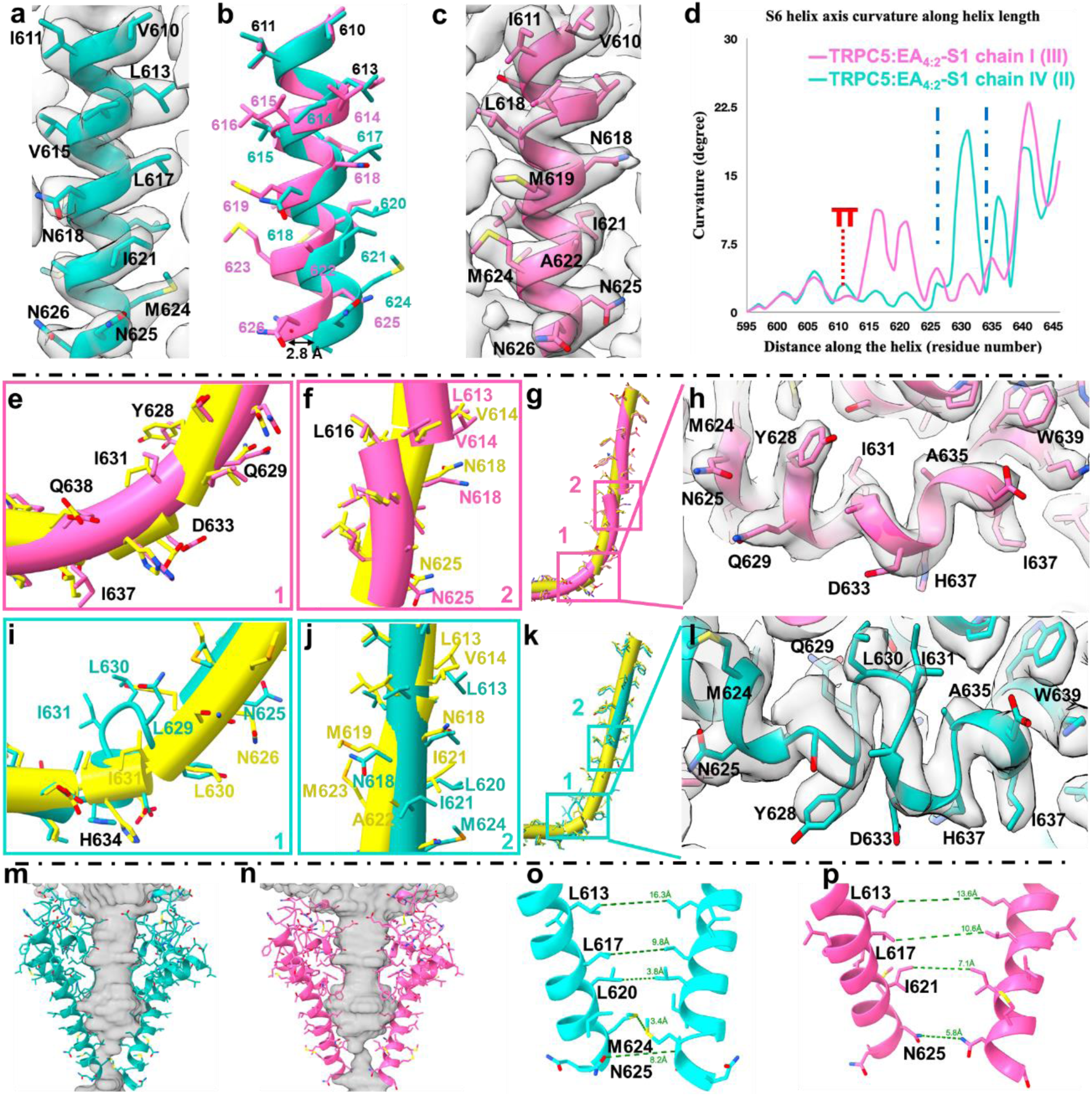
π-bulge to α-helix transitions in two opposing TRPC5 subunits result in a symmetry break in the channel pore. a-c, Cryo-EM map and model of S6 from two adjacent monomers showing a full transition of π-bulge to α-helix in TRPC5 subunit IV (cyan; a) and a partial shift in π-bulge position in TRPC5 subunit I (pink; c), with model superimposition illustrating the change in helix angles after of π-bulge forming residues (b). **d,** Curvature analysis of S6 helix from TRPC5 subunits I (pink) and IV (cyan). The position of Val610 (the start of the π-bulge) is indicated in red. **e-l,** Comparative analysis of the S6 helix from TRPC5 subunits I (pink) and IV (cyan) in TRPC5:EA**_4:2_**-S1 with the S6 helix from TRPC5:EA^PA^-S1 (yellow). Close-ups of main curvature points (e,i,g,k), π-helix (f,j) and the TRP helix-S6 linkage loop (h,l). The cryo-EM densities and models of TRPC5 subunits I (pink) and IV (cyan) in the connecting region between TRP helix and S6 show the formation of a finger-like loop in TRPC5 subunit IV (h,l). **m-p,** Pore shape analysis showing a transition of C4 to C2 symmetry after the TRPC5 selectivity filter/π-bulge forming residues.

Exploration of the EA binding sites in TRPC5:EA**_4:2_**-S1 revealed an interaction pattern similar to that found in TRPC5:EA^PA^-S1, involving Gln573(I), Leu528(I), Tyr524(I), Phe520(I), Leu521(I) and Val610(IV) (**Figure 6e**). When comparing sites occupied by EA to those occupied by lipid, clear differences were observed between conformations of the different Phe520 residues and their aromatic interaction networks (**Figure 6d,f**). In pockets occupied by EA, Phe520 occupies a position similar to that in TRPC5:EA^PA^-S1, i.e. pointing upwards towards the symmetry axis (**Figure 6g-i**). This facilitates the formation of a triangular aromatic network with Tyr524 and the phenyl ring of EA’s 6-cinnamate substituent, while C-π interactions between Leu620 and Tyr524 further stabilise this conformation. Surprisingly, in pockets occupied by lipid, Phe520 does not adopt the conformation seen in the ‘apo’ structure TRPC5^PA^. Instead, it resembles the conformation found in TRPC5:EA**_4:4_**-S1, i.e. pointing downwards, with rearrangement of the S6 helix allowing a S-π interaction with Met619 and a C-π interaction with Leu506 (**Figure 6d**). Superimposition of the region around Phe520 of subunits I/III and II/IV reveals clear conformational differences (**Figure 6g-i**). For S5, the local RMSD – including five residues above and below Phe520 – is 2.1 Å. A similar analysis of S6, covering 5 residues above and below Leu620, reveals an even larger RMSD of 5.1 Å. These changes around Phe520 suggest that binding of two EA molecules modifies the energetic landscape of the S5 helices in all four TRPC5 subunits through a direct interaction that results in the formation of aromatic-aromatic bridges within subunits I and III, and allosteric rearrangement of the S6 helices of subunits II and IV.

Expansion of the RMSD analysis to include the entire S5 and S6 helices showed deviations of 1.5 Å and 4.1 Å, respectively. When examining the curvature of the S6 helix, we found that, in adjacent subunits (i.e. I/III and II/IV), residues overlap well until Leu613, which is responsible for the formation of a π-bulge in ‘apo’ TRPC5 structures (e.g. **Extended Data Figure 7**). In subunits I/III, the π-bulge changes its side and position, moving from Leu613-Val614 to Val615-Leu616. In subunits II/IV, we see a transition from a π-bulge to an α-helix (**Figure 7a-d**). As in TRPC5:EA**_4:4_**-S1, this transition is triggered by a rotamer change of Leu613, leading to rotation of the S6 helix, shifting each residue by one position. The disappearance of the S6 π-bulge of subunits I/III causes the S6 helices to adopt a straight conformation until Asn626. In contrast, the positional shift in subunits II/IV results in the formation of kinked helices between Leu613-Trp639. The differences between the S5/S6 helices from different subunits of TRPC5:EA**_4:2_**-S1 and those of TRPC5:EA^PA^-S1 are even more striking (**Figure 7e-g,i-k**). The S5/S6 helices of TRPC5:EA**_4:2_**-S1 adopt unique conformations. This includes the surprising formation of a finger-like loop between residues Ser627 and Ala635 of subunits II and IV, leading to re-adjustment of the symmetry and position of the corresponding TRP helices compared to subunits I and III (RMSD 0.9 Å) (**Figure 7h,l**). A similar difference was noted when comparing the finger-like loop to the corresponding domain in TRPC5:EA^PA^-S1 (**Figure 7i**). The finger-like loop forms immediately after the channel gate, resulting from the flip of Tyr628, and extends upstream of the S6 helix that connects the loop.

Pore analysis of TRPC5:EA**_4:2_**-S1 reveals a break in symmetry in the pore after Leu613 (**Figure 7m-p; Extended Data Figure 11**). Up to this point, a pseudo-C4 symmetric pore shape can be observed, with similar distances between residues in opposing subunits (**Figure 7m-p**). After Leu613, a transition to C2 symmetry can be seen in the lower section of the pore (**Figure 7m,n**). Between subunits I and III, a slight expansion can be seen (**Figure 7p**), as determined by the side shift of the π-bulge and the formation of strongly kinked S6 helices (**Figure 7f,g**). The transition to an α-helix in subunits II and IV creates a double hydrophobic gate between residues Leu620 and Met624 (**Figure 7o**).

In comparison to TRPC5:EA**_4:4_**-S1, where a similar transition occurs, the binding of two EA molecules per TRPC5 tetramer in TRPC5:EA**_4:2_**-S1 leads to the repositioning of the S6 segment, narrowing the distance between subunits II and IV (**Figure 7m,o**). The main bottleneck is formed between residues Met624(II) and Met624(IV), which have side chains pointing towards the centre of the pore. In contrast, in TRPC5:EA**_4:4_**-S1, the main bottleneck is located at the level of Leu620, with the side chains of Met624 directed downwards and outwards from the symmetry axis. Measuring the distances between the pore-forming residues in subunits II and IV of TRPC5:EA**_4:2_**-S1 reveals significant differences compared to TRPC5:EA**_4:4_**-S1, particularly at the levels of Leu620, Met624, and Asn625 (cf. **Figure 5j** and **Figure 7o**). This suggests an intermediate conformation. It is likely that the channel requires full occupancy by EA (four EA molecules per TRPC5 tetramer) for a complete transition to occur. Interestingly, analysis of the pores of structures of TRPC5 with mixed EA occupancy (TRPC5:EA**_mix_**-S1, TRPC5:EA**_mix1_**-S2 and TRPC5:EA**_mix2_**-S2, containing 1-3 EA molecules per tetramer) suggests that transition of the S6 π-bulge to an α-helix in a single subunit could be sufficient to allow a water molecule to pass (**Extended Data Figure 11g-o**). We therefore hypothesise that different EA binding stoichiometries could result in different conductive states of TRPC5 channels.

## Discussion

Many TRP ion channels are modulated by plant-derived natural products, suggesting evolutionary relationships between TRP proteins expressed in various species and the metabolites found in their dietary sources. The guaiane sesquiterpene derivative EA is an intriguing example of a TRP channel-modulating plant metabolite, which activates TRPC1/4/5 channels with high potency, efficacy and selectivity.

Our study provides the first structural insight into the interaction between the plant natural product EA and TRPC5, revealing mechanisms of TRPC5 ligand recognition, channel dynamics and channel gating. Our TRPC5 structures, supported by functional evaluation of TRPC5 variants, reveal that EA binds to a conserved lipid binding site between TRPC5 subunits, similar to xanthine-based TRPC1/4/5 modulators such as the inhibitors Pico145^39^ and HC-070^40^, the latter of which is the subject of multiple clinical trials. We also show the importance of the aromatic interaction network around Phe520 of TRPC5 with conserved methionine residues. This network is crucial to channel structure and function, including its ability to respond to voltage and EA. By varying the method of ligand delivery and using 3DVA, we determined structures of TRPC5 with different EA binding stoichiometries, revealing key EA-induced conformational changes of membrane helices and residues. Together, these findings will be crucial to structure-based design of TRPC5 modulators and to understanding the pharmacological effects of TRPC5 inhibitors in studies that use EA as an activator.

The finding that EA binds to the same site of TRPC5 as xanthine-based modulators is consistent with the pharmacological profile of this compound class. For example, Pico145 shows competitive inhibition of TRPC4/5 channels with respect to EA^46,47^ and AM237, Pico145-DA and Pico145-DAAlk are partial agonists of TRPC5 that suppress EA responses of TRPC5, and inhibit EA responses of other TRPC1/4/5 channels^51,62^. Analysis of the EA binding site also explains the importance of EA’s glycolate substituent^6^, which forms a dynamic hydrogen bond network with Trp577, Gln573 and Arg557. In vivo, EA’s glycolate ester bond is hydrolysed to produce the main EA metabolite, (-)-englerin B (EB)^6,29^, which results in a dramatic loss of activity as a TRPC1/4/5 activator. This is also consistent with medicinal chemistry efforts, which reveal that changes to the EA glycolate group mostly result in loss of efficacy or potency at TRPC4/5 channels^63^. It is noteworthy that the EA analogue A54, which contains an ether bond in place of EA’s glycolate ester bond, does not activate TRPC1/4/5 channels, but is a competitive inhibitor of the channels’ EA response, while potentiating the effect of other agonist such as Gd^3+^ or sphingosine-1-phosphate^48^.

A study examining the agonistic effects of EA on TRPC5 variants suggested that three specific charged residues (Lys554, His594, and Glu598) are directly involved in EA binding^44^. In contrast, our structures reveal that these residues are located relatively far from the actual binding site, and do not interact directly with EA. We expect that these residues exert an allosteric effect on EA modulation, likely by disrupting hydrogen bond formation between the hydroxyl group of EA and proximal residues Gln573 (as in TRPC5:EA^PA^-S1, TRPC5:EA^PA^-S2, TRPC5:EA_4:4_-S1 and TRPC5:EA**_4:2_**-S1) or Arg557 (as in TRPC5:EA**_4:4_**-S1).

Given the high degree of sequence conservation between TRPC1, TRPC4 and TRPC5 – especially in the EA binding site (**Extended Data Figure 5e**) – and the ability of EA to activate homo-and heteromeric TRPC1/4/5 channels, we expect that our results extrapolate to the entire subgroup. The residues interacting with EA are fully conserved between TRPC4 and TRPC5, consistent with the high potency and efficacy on both TRPC4 and TRPC5 channels. While TRPC1 may not form functional homomeric channels, it forms heteromeric channels with TRPC4 and TRPC5^64–67^. Comparison of TRPC5 residues found to interact with EA with those from TRPC1 reveal a less conserved pattern. We speculate that some sequence differences may have limited effects; for example, Tyr524 is a phenylalanine in TRPC1, but EA retains its efficacy against TRPC5_Y524F_. However, Arg557 and Gln573 are phenylalanines in TRPC1, which could prevent the formation of a hydrogen bond network with the hydroxyl group EA’s glycolate substituent. Additionally, the substitution of Val614 in TRPC5 by an isoleucine (present in TRPC1) may cause a steric clash with the phenyl ring of EA’s 6-cinnamate substituent. These differences may prevent binding of EA to TRPC1-adjacent binding sites. Nevertheless, previous studies have shown that EA exhibits similar potency in both homo-and heteromeric TRPC1/4/5 channels, with EC_50_ values in the low nM range^5,6^. How can these observations be explained?

The exact composition and stoichiometry of TRPC1/4/5 heteromers, as well as their ability to display different biophysical properties, is only partially understood. Two recent publications – a proteomics-based interactome study of TRPCs in mouse brain^67^ and a cryo-EM analysis of the TRPC1/C4 heteromer^37^ – suggest asymmetric assemblies of TRPC4/5 with TRPC1 in a 3:1 stoichiometry. This suggests that one TRPC1 subunit can influence the properties of the entire channel, and that partial occupancy (i.e. 1-3 molecules of EA per tetramer) may be sufficient to activate heteromeric TRPC1/4/5 channels. According to our structural analysis, this would likely result in asymmetric channel pores, which could underlie the reduced calcium permeability, preferential permeability of monovalent cations, and distinct voltage-current relationship of heteromeric TRPC1/4/5 channels.^37,64,65,68–70^

Why does EA activate TRPC1/4/5 channels but not TRPC3/6/7 channels? In the structures of TRPC3 (PDB 7DXB) and TRPC6 (PDB 7DXF),^71^ the loop between the S4 segment and the pore helix is shorter, and the positions of TRPC5 Ala602 and Gln573 are occupied by a tyrosine and a lysine, respectively. This change may prevent formation of a hydrogen bond network with EA’s glycolate. Additionally, substituting Leu572 of TRPC5 with a phenylalanine (as in TRPC3/6/7) would lead to a significant steric clash, requiring substantial rearrangements to fit an EA molecule. In addition, replacement of two adjacent phenylalanines (Phe599 and Phe569) flanking the EA binding site of TRPC5 by polar or charged residues (asparagine and aspartate, respectively) in TRPC3/6/7 creates a distinctly different environment in the equivalent lipid binding site of TRPC3/6.

Recent studies show that Met-aromatic motifs have both structural and functional roles in both soluble proteins and membrane proteins, especially ion channels. For example, in SUMO, a conserved Met-Phe pair is critical for its β-grasp fold, facilitating noncovalent interactions with its ligands. It is well-established that Ile-Phe-Met motifs are involved in the fast inactivation of Na_v_ channels and their ability to sense voltage^72,73^. Furthermore, recent studies have shown that Phe-Met motif can act as latches in the activation of Orai1 and play a crucial role in the voltage-sensing capability of HCN1^74^. Notably, propofol has been able to rescue this function in disease-relevant mutants by mimicking the Met-aromatic interaction, further underscoring the critical functional role of these motifs. We show that a Met-aromatic motif (Met623-Phe520) is vital for both voltage-and EA-mediated modulation/activation of TRPC5. Alanine substituting either of these residues strongly reduces EA-and voltage-evoked currents. Interestingly, replacing Phe520 with a different aromatic residue alters the voltage sensitivity and channel kinetics in a contrasting way, with TRPC5_F520W_ resulting in a gain-of-function but TRPC5_F520Y_ leading to a loss of its voltage-sensing ability. The reason for the loss of voltage sensing by TRPC5_F520Y_ remains unclear; we hypothesise that the different ring electronics prevent the formation of the Met-aromatic interaction. These findings suggest that Phe520 plays a switch-like role in TRPC5 channel modulation. While TRP(C) channels other than TRPC1/4/5 are insensitive to EA, the conservation of residues in the aromatic interaction network of Phe520 in TRPC5 implies that similar phenylalanine switches may be important for the functionality of other TRP(C) channels.

Our cryo-EM structures highlight the dynamic nature of TRPC5 by revealing transitional conformations between closed and open structures. Intriguingly, we demonstrate that it is not necessary for the channel to have full occupancy in order to induce significant conformational changes. However, it remains unclear whether partial occupancy is sufficient to fully open the channel. Our findings suggest that varying stoichiometries of EA binding could result in different states with potentially unique conduction properties. These insights may help explain the conundrum of TRPC polymodal signal integration alongside the different biophysical properties of TRPC1/4/5 heteromers (see above).

The finger-like loop between residues Ser627 and Ala635 observed in TRPC5:EA**_4:2_**-S1 comprises a functionally important Asp633 that determines the double rectification of the current-voltage (I/V) relationship typical of TRPC4/5 (but also of TRPC3/6/7) by electrostatically attracting intracellular Mg^2+^.^75^ Our comparison of the average I/V curves measured at the maximum of the peak of individual EA responses reveals that the inflection differs among the different TRPC5 variants (**Figure 2d-j**; **Figure 5n**; **Extended Data Figure 4**). Some of the constructs, such as TRPC5_Y524F_, TRPC5_Y524W_ and TRPC5_L613A_, exhibit a less flat part at positive voltages than others (e.g., TRPC5_F520W_, TRPC5_L521A_, TRPC5_L620F_). Because all these mutations affect the ability of the channel to translate EA binding to channel opening (gating), we speculate that conformational changes in the finger-like loop region, which we found to be dependent on the degree of EA occupancy, may be responsible for the changes in the shape of the TRPC5 current-voltage relationship.

While we have gained valuable structural insights into channel dynamics, we did not find conducting states of TRPC5. This is likely the result of a combination of factors: the low open probability of TRPC5, the absence of a voltage gradient, and the replacement of the native membrane environment (likely incorporating lipids critical to channel modulation) by a belt of amphipathic polymer. Therefore, we are cautious about proposing a full gating mechanism. Further research is necessary to improve our understanding of the gating and modulation mechanisms related to TRPC1, TRPC4, and TRPC5.

In summary, our new TRPC5 structures have uncovered the EA binding pocket in high resolution. Furthermore, the various conformational states presented here extend our understanding of the TRPC5 dynamic transition between the closed and open states. Moreover, we show for the first time that a Met-aromatic motif can play a switch-like role in TRPC5, a concept that likely extends to other TRP(C) channels. Finally, we consider that the different high-resolution structures determined in this study, alongside the identified functional role of the aromatic networks, are an excellent starting point for structure-guided drug discovery projects.

## Materials and Methods

### Chemicals

(-)-Englerin A (EA; PhytoLab) was stored at –80 °C as a 10 mM or 1 mM solution in DMSO. Where stated, 0.01% pluronic F127 (PA; Sigma-Aldrich) was included in experimental solutions of EA. Fura-2 AM (ThermoFisher) was stored at –20 °C as a 1 mM solution in DMSO. AM237 was prepared according to published procedures^51^. BTD (Tocris) and AM237 were stored at-20 °C in DMSO at 100 mM (BTD) or 10 mM (AM237) stock. Carbamoylcholine chloride (carbachol; Cayman Chemical) and gadolinium(III) chloride hexahydrate (Gd^3+^; Fluorochem) were diluted in SBS to 100 mM on the day of the experiment.

### Production of purified TRPC5 protein

#### Plasmid

Production of TRPC5 protein was performed using the previously described construct MBP-hTRPC5_Δ766–975_, which contains human TRPC5 in C-terminally truncated form (Δ766–975) with an N-terminal maltose-binding protein tag followed by a PreScission protease cleavage site^39^. Bacmids and baculoviruses were produced according to the Bac-to-Bac protocol (Invitrogen). The BacMam vector^76^ was a kind gift from Professor Eric Gouaux (Vollum Institute).

#### Protein expression and purification

P2 virus was added to Freestyle™ 293-F Cells (ThemoFisher Scientific) at 2.0 million cells per ml, so the virus was at a final concentration of 10% v/v. The cells were kept in Gibco FreeStyle 293 Expression Medium (Invitrogen), shaking, at 37 °C and 5% CO_2_. After 16-24 h, 5 mM sodium butyrate (Sigma Aldrich) was added, and the temperature was lowered to 30 °C. After a further 40 h, cells were harvested by centrifugation, washed with 30 ml PBS and stored at-20 °C. The protein purification protocol was adapted from Duan et al.^38^, unless stated otherwise. All detergents (and amphipol: PMALC8) were supplied by Generon. For **TRPC5^PA^**, **TRPC5:EA^PA^-S1** and **TRPC5:EA^PA^-S2** sample preparation, a 200 ml cell pellet was thawed and resuspended in 20 ml of buffer containing 1% DDM, 0.1% CHS, 150 mM NaCl (Sigma Aldrich), 30 mM HEPES (Sigma Aldrich) pH 7.4, 1 mM DTT (Fisher Scientific Ltd), and protease inhibitor cocktail (Sigma Aldrich), and was then incubated while rotating at 4 °C for 1 h. The insoluble material was removed by centrifugation at 10,000 × g for 1 h at 4 °C. The soluble fraction was incubated with 1 ml bed volume of pre-washed amylose resin (New England Biolabs) for 12-16 h, rotating at 4 °C. The resin was washed with 50 ml of buffer containing 0.1% DDM and 0.01% CHS, 150 mM NaCl, 30 mM HEPES pH 7.4, and 1 mM DTT. The resin was resuspended in 4 ml of buffer containing 0.2% PMAL-C8, 150 mM NaCl, 30 mM HEPES pH 7.4, 1 mM DTT and rotated for 2 h at 4 °C followed by addition of another 4 ml of the same buffer, and incubated for an additional 4 h at 4 °C. The detergent was removed by the addition of Bio-Beads (Bio-Rad) at a ratio of 10 mg/ml and incubated for 16-20 h. The resin was washed with 20 ml of buffer without DDM (150 mM NaCl, 30 mM HEPES pH 7.4, 1 mM DTT). The sample was eluted in 2 ml of the same buffer supplemented with 50 mM maltose (Sigma Aldrich). The eluate was subjected to centrifugation at 20,000 × g for 15 min at 4 °C to remove any precipitated material. The supernatant was concentrated stepwise to 0.8-1 mg/ml with 100 kDa cut-off Vivaspin 500 concentrators (Sigma Aldrich). For **TRPC5_EA_** sample, purification was varied as above with a few modifications. Namely, after solubilisation, EA was added to the soluble fraction to a final concentration of 100 μM. Further, all buffers were supplemented with EA at the same concentration, resulting a constant concentration of 100 μM EA over the entire purification process.

### Cryo-EM studies

#### Grid preparation and data collection

For cryo-EM studies leading to structures TRPC5^PA^, TRPC5:EA^PA^-S1 and TRPC5:EA^PA^-S2, we incubated purified TRPC5 in PMAL-C8 at 1.0 mg/ml with 100 µM of EA, previously diluted in protein buffer with 10% DMSO and 1% PA (EA was taken from a 10 mM DMSO stock, while PA was added from a 10% stock, never exceeding 1% DMSO final concentration in protein samples).

For TRPC5^PA^, a 3.5 µl aliquot of the sample was applied to an UltrAuFoil Holey R1.2/1.3, 300 mesh grid (Quantifoil), which had been glow-discharged twice for 45 s using a Pelco easyGlow glow discharge unit. In case of TRPC5:EA^PA^, a 3 µl aliquot was applied to a HexAuFoil grid (Quantifoil) that was glow discharged twice, first time for 90 s followed by additional 45 s using a Pelco easyGlow glow discharge unit. The grids were vitrified using an FEI Vitrobot IV, at 100% humidity and 4 °C. In case of TRPC5^PA^, the samples were vitrified with a blot time of 6 s and blot force 1, while in case of TRPC5:EA^PA^, the blotting time was increased to 10 s and the blotting force to 6. The grids were loaded into an FEI Titan Krios transmission electron microscope (Astbury Biostructure Laboratory, University of Leeds) operating at 300 kV, fitted with a Falcon 4i direct electron detector and Selectris. Automated data collection was carried out using EPU software, with fringe-free imaging in counting mode, using a defocus range between −0.7 to −3 µm in 0.3 µm increments. We collected a total of 5,000 EER movies for TRPC5^PA^ at a nominal magnification of 165k resulting a pixel size of 0.74 Å and a total dose of ∼35 e−/Å2. In case of TRPC5:EA^PA^, we collected 12,263 movies, at a nominal magnification of 270k, with a pixel size of 0.46 Å and a total dose of ∼40 e−/Å2. In all cases the fractions were combined to give an exposure of 1 e−/Å2 per frame.

In case of **TRPC5:EA** (i.e. no PA added), 3.5 µl aliquot of the concentrated sample (already containing EA, with no further incubation) was applied to an UltrAuFoil Holey R1.2/1.3, 300 mesh grid (Quantifoil) which had been glow-discharged twice for 45 s using a Pelco easyGlow glow discharge unit. The sample was vitrified using the same device as above, using a blotting time of 6 s and blotting force 6. We collected data on a similar microscope to previously described, but without the Selectris. EPU software was used to set up automated data collection, with fringe-free imaging in counting mode, and a defocus range between −0.7 to −3 µm in 0.3 µm increments. We acquired 15,538 EER movies, at a nominal magnification of 130k, with a pixel size of 0.82 Å and a total dose of ∼35 e−/Å2. As with the other data sets, the fractions were combined to give an exposure of 1 e−/Å2 per frame.

#### Image processing

Overviews of the image processing protocols are shown in **Extended Data Figure 2** and **Extended Data Figure 8**. All processing was completed in CryoSPARC^77^ (v4.2 or v4.4) unless stated otherwise. The initial drift and beam-induced motions were corrected for using *Patch Motion Correction* while CTF estimation was performed using *Patch CTF Estimation*, both with default settings. We used a TRPC5 map, previously obtained in-house, down-filtered to 20 Å, to generate 50 templates using *Create templates* tool in CryoSPARC, which were further used for template-based particle picking. The pre-processing part was identical for all three samples.

For the **TRPC5^PA^** sample this resulted, after filtering the picks based on the NCC score (>0.25), in a stack of ∼1.2 million particles which were binned 4 times. After several rounds of 2D classification, we curated the particle stack to 120k which was further used to generate five initial models using ab initio reconstruction with standard parameters. From the obtained models, at least one had the expected shape. The initial models were used to confirm the good map and further sort the particle stack using *Heterogeneous Refinement* with default setting. Further, the whole particle stack was put through iterative rounds of heterogeneous refinement and 2D classification, and when all noisy classes were removed, the particles were re-extracted with the full box-size. Using this approach, we observed that our final stack of good particles increased by roughly 20% while also improving particle orientation by either increasing the number of rare orientation or by increasing the particle number in low-populated 2D classes. After another two iterative rounds of heterogeneous refinement, we ended up with a stack of 125k good particle. A final round of 2D classification was performed to remove classes that did not show high resolution features. The final particle stack (∼225k) was run in a non-uniform refinement, with C4 symmetry, defocus refinement, CTF refinement enabled. Further we performed reference-based motion correction followed by another round of *Non-Uniform Refinement*^78^ with 5 extra final passes, and having defocus refinement, CTF refinement (all option true), anisotropic magnification and EWS correction active and imposing a C4 symmetry. The final map had a global resolution of 2.38 Å as estimated based on the gold standard FSC = 0.143 criterion. We used ResolveCryoEM^79^ features of Phenix^80^ to improve the interpretability of the map.

Similar image processing workflows were followed for TRPC5:EA^PA^ and TRPC5:EA, but with a few modifications.

In case of **TRPC5:EA^PA^**, after filtering the picks based on the NCC score (>0.25), we ended up with a stack of ∼896k particles, which were extracted and binned 4 times. Iterative rounds of 2D classification were used to remove bad classes, and ∼97k particles were used to generate 6 *ab-initio* classes. These classes were subjected to heterogeneous refinement, resulting in a best class with ∼47k particles. The full particle stack was then put through this *Heterogeneous Refinement*, resulting in ∼187k particles in the good class. 2D classification was used to further clean the particle stack. A final round of 2D classification was performed to remove classes that did not show high resolution features. The final particle stack (∼110k) was run in *Non-Uniform Refinement*, with C4 symmetry, defocus refinement, CTF refinement enabled. Further we performed reference-based motion correction followed by another round of non-uniform refinement with 5 extra final passes, and having defocus refinement, CTF refinement (all option true) and EWS correction active and imposing C4 symmetry. This resulted with a map with a global resolution of 2.4 Å. Following visual inspection of the map, we noticed several regions with high flexibility, especially in the ankyrin domain region. Therefore, the data were subjected to 3D variability analysis (3DVA)^81^, with 3 clusters, filtered to 3.5 Å. The two clusters representing protein (making up 34% of the particles-state 1, and 63% of the particles – state 2) were subjected to non-uniform refinement, with C4 symmetry, defocus refinement, CTF refinement. Following reference motion correction, a final round of non-uniform refinement was performed with 5 extra final passes, and having defocus refinement, CTF refinement (all option true), anisotropic magnification and EWS correction active while also imposing C4 symmetry. These resulted with maps at final global resolutions of 2.49 Å (state 1), and 2.54 Å (state 2) as estimated based on the gold standard FSC = 0.143 criterion. We used ResolveCryoEM^79^ features of Phenix^80^ to improve the interpretability of the map.

In case of **TRPC5:EA** (i.e. no PA added), after filtering the picks based on the NCC score (>0.25), we ended up with a stack of 4.5 million particles, which were extracted and binned 4 times. Iterative rounds of 2D classification were used to remove bad classes, and ∼250k particles were used to generate 5 ab initio classes. These classes were subjected to heterogeneous refinement, resulting in a ‘best class’ with ∼180k particles. The full particle stack was then put through *Heterogeneous Refinement*, resulting in ∼300k particles in the good class. A final round of 2D classification was performed to remove classes that did not show high resolution features. The final particle stack (∼257k) was run in *Non-Uniform Refinement*, with C4 symmetry, defocus refinement, CTF refinement enabled. Further we performed reference-based motion correction followed by another round of non-uniform refinement with 5 extra final passes, and having defocus refinement, CTF refinement (all option true) and EWS correction active and imposing C4 symmetry. This resulted with a map with a resolution of 2.6 Å. Following visual inspection of the map, we noticed several regions with high flexibility, especially in the ankyrin domain region, xanthine binding site and pore gate region. Therefore, the data were subjected to 3D variability analysis with 3 components. After several trials, we obtained the best separation using six clusters, 3 clusters for each state. Each cluster was then refined using non-uniform refinement, using C4 relaxed symmetry, with defocus and CTF refinement. Following reference motion correction, a final round of non-uniform refinement using C4 relaxed symmetry was performed with 5 extra final passes, and having defocus refinement, CTF refinement (all option true), anisotropic magnification and EWS correction active. Maps representing state 1 represented approximatively 30 % of the data and were split based on EA stoichiometry as follows: 4 EA bound (6.5% with a final resolution of 2.83 Å), 2 EA bound (14.9% and a final resolution of 3.04 Å), mixed EA occupancy (7.2% with a final resolution of 3.22 Å). State 2 classes represented approximative 70% and were split based on EA stoichiometry in: a class with 2 EA bound (22.2% with a final resolution of 2.97 Å), and two classes with mixed EA occupancy (22.2% and 24.2% with final resolutions of 3.15 Å and 3.23 Å, respectively. We used ResolveCryoEM^79^ features of Phenix^80^ to improve the interpretability of the map

#### Model building

The models for all structures were built using ModelAngelo^82^. The models obtained were inspected and manually completed in Coot^83^. Several rounds of real-space refinement were performed in Phenix^80^ before fitting the ligand. We used Coot to manually fit the small molecule EA, generated with the restraints from AceDRG program^84^ in the CCP-EM (v1) software package^85^, starting from a SMILE string. After ligand fitting, we manually checked the structures in Coot and performed additional real-space refinement in Phenix. Protein-ligand interactions were visualised with the PLIP web tool (https://plip-tool.biotec.tu-dresden.de/plip-web/plip/index)^86^ and ChimeraX^87^. All structural images were produced using ChimeraX (v1.8) and PyMOL (v2.6 LTS) (Schrödinger, LLC. 2010). Aromatic interactions were identified using the Arpeggio web server (https://biosig.lab.uq.edu.au/arpeggioweb/)^59^. Pore shape and radius was calculated using a local installation of PoreAnalyser (https://poreanalyser.bioch.ox.ac.uk)^60^. Helix curvature was calculated using the HELANAL-PLUS web server (http://nucleix.mbu.iisc.ac.in/cgi-bin/helanalplus/helanalplus.cgi)^54^ and the Bendix program (https://sbcb.bioch.ox.ac.uk/Bendix/)^88^ inside the VMD (v.1.9.4a57) software package^89^.

#### Generation of TRPC5 constructs

TRPC5 constructs for patch clamp electrophysiology were generated by introduction of point mutations into hTRPC5 in the pCMV6-XL5 vector (OriGene Technologies) using QuikChange II XL Site-Directed Mutagenesis Kit (Agilent Technologies). Primers are listed in Table 1. Mutations were confirmed by DNA sequencing (Eurofins Genomics).

**Table 1.**
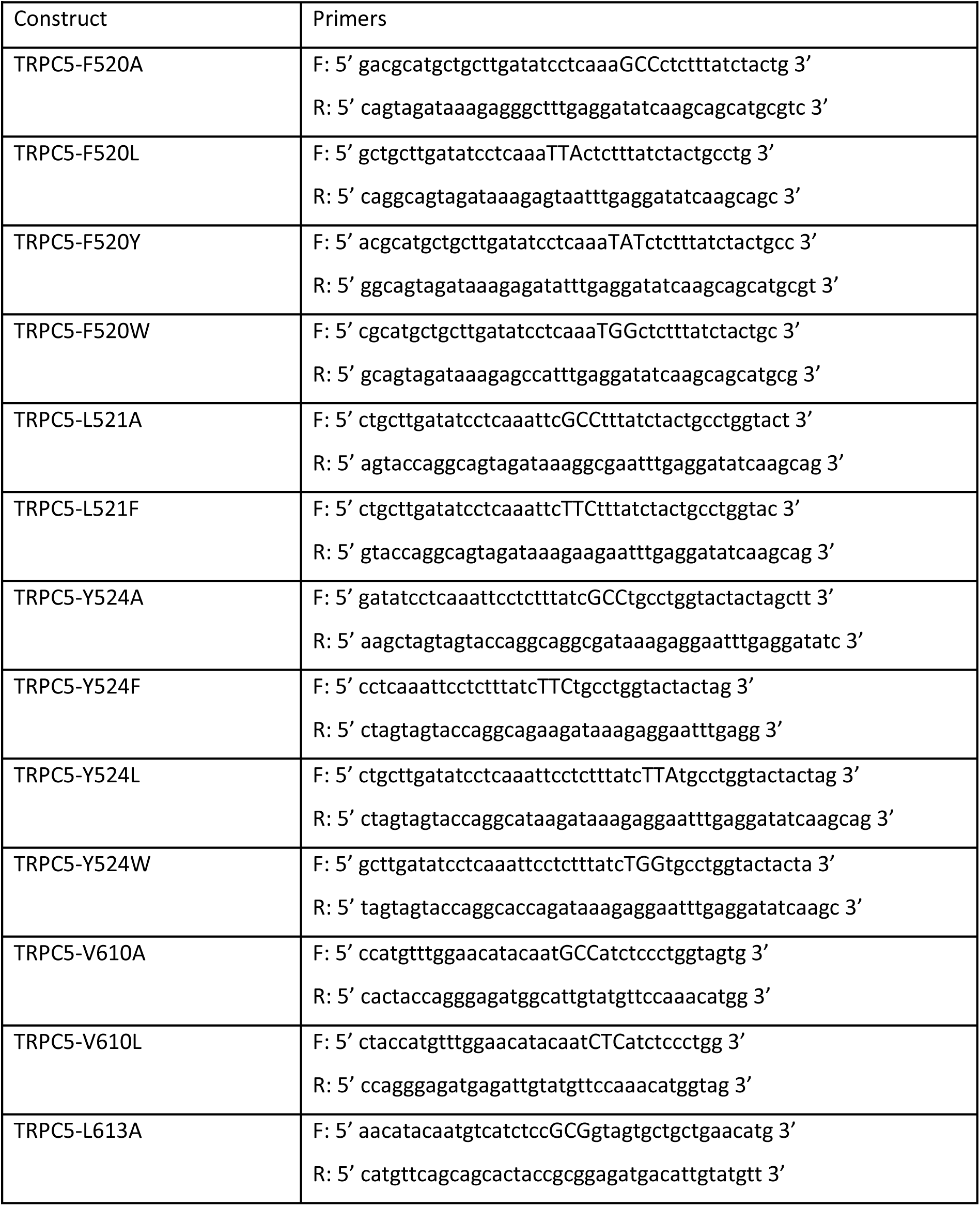
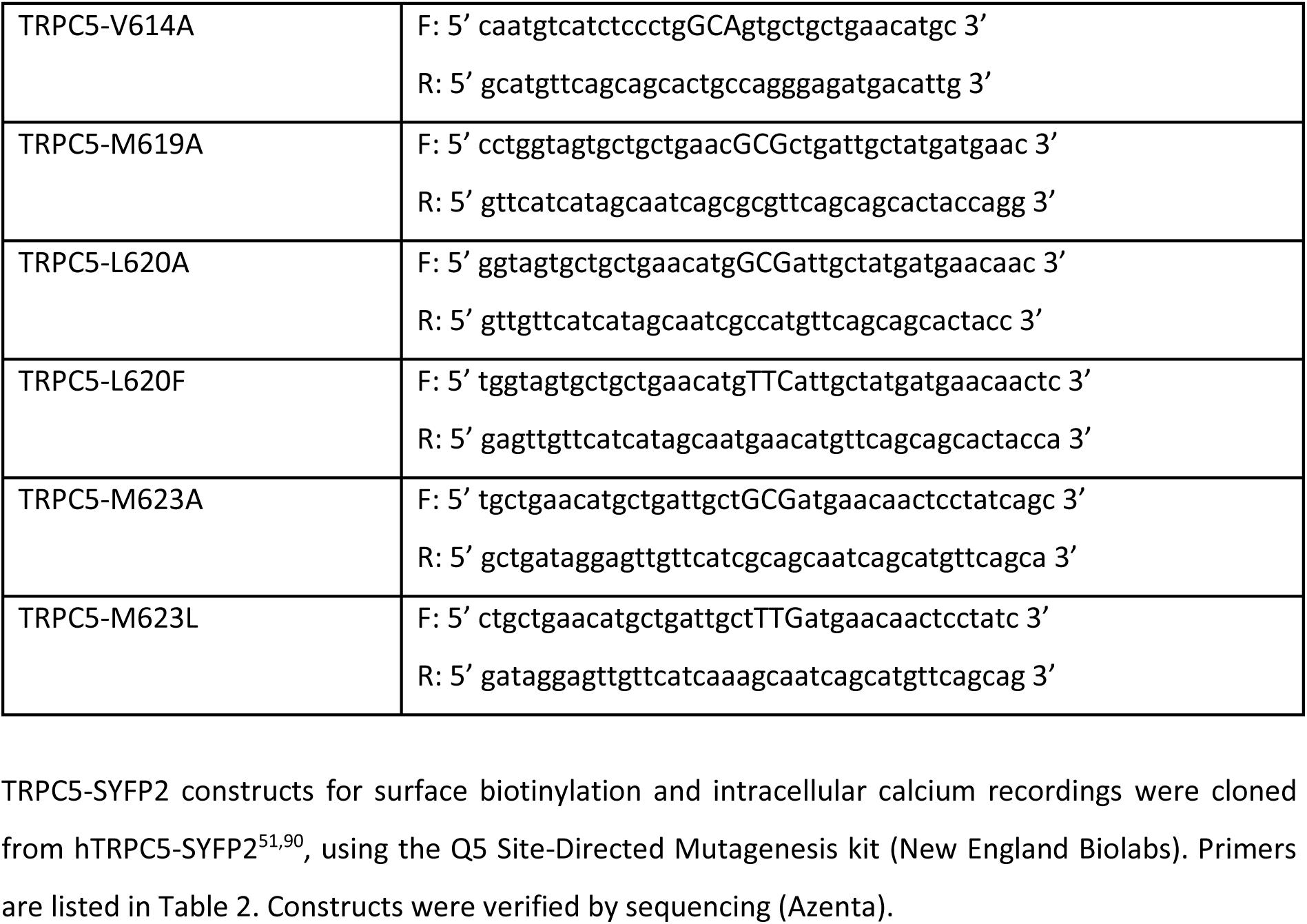
Primers for TRPC5 constructs.

#### Primers for TRPC5 constructs for patch-clamp electrophysiology measurements

TRPC5-SYFP2 constructs for surface biotinylation and intracellular calcium recordings were cloned from hTRPC5-SYFP2^51,90^, using the Q5 Site-Directed Mutagenesis kit (New England Biolabs). Primers are listed in Table 2. Constructs were verified by sequencing (Azenta).

**Table 2.**
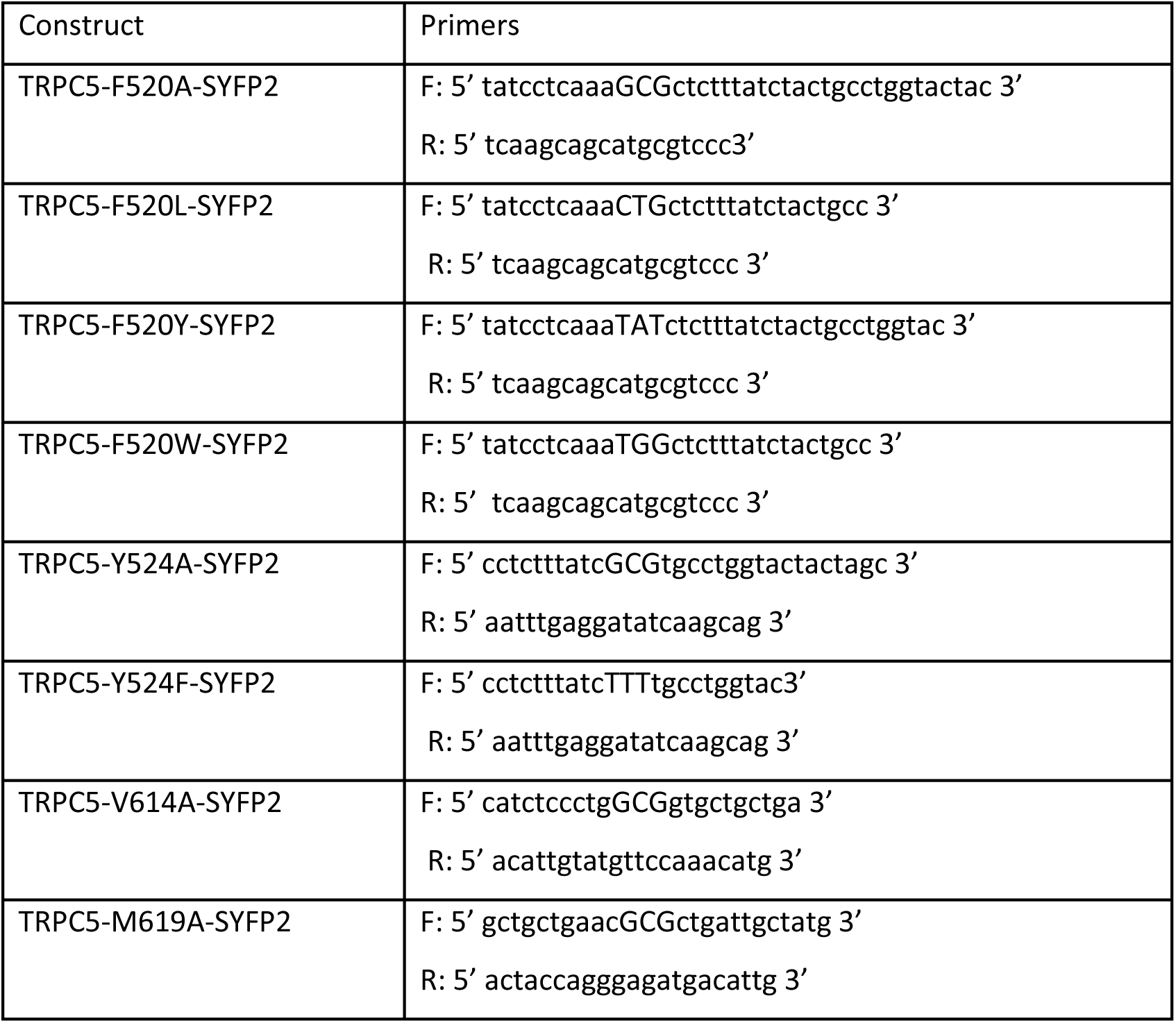
Primers for TRPC5-SYFP2 constructs.

### Functional evaluation of TRPC5 variants

#### Cell culture and transfection

Human embryonic kidney 293T cells (HEK293T, ATCC) were cultured in Opti-MEM I medium (Thermo Fisher Scientific) supplemented with 5% fetal bovine serum (PAN-Biotech). Before transfection, cells were placed into 24-well plates coated with poly-L-lysine and collagen. After reaching ∼70% confluency, cells were transiently co-transfected with 200 ng of eGFP plasmid in the pcDNA3.1 vector (a kind gift from Jan Teisinger; Institute of Physiology, Prague) and with 300 ng of plasmid encoding wild-type or mutant TRPC5 construct using the magnet-assisted transfection technique (PolyMag Neo, OZ Biosciences). Cells were then plated on poly-L-lysine-coated glass coverslips. Electrophysiological recordings were performed 24-48 h after transfection. At least two independent transfections were used for each experimental group. The wild-type TRPC5 channel was regularly tested alongside these experiments.

#### Patch-clamp electrophysiology

Whole-cell membrane currents were recorded with an Axopatch 200B amplifier and the software pCLAMP 10.6 (Molecular Devices). Data were filtered at 2 kHz using the low-pass built-in 8-pole Bessel filter and digitised at 5-10 kHz using the Digidata 1550B analog-to-digital converter controlled by Clampex 10.6 (Molecular Devices). Patch pipettes were prepared from borosilicate glass capillaries with 1.5-mm outer diameter (Science Products) pulled on a horizontal puller P-87 (Sutter Instrument) and heat-polished with microforge MF-83 (Narishige) to a final resistance 3-5 MΩ. Two voltage stimulation protocols were used; 100 ms voltage steps from-80 to +200 mV (with 20 mV increments) applied from a holding potential of-70 mV and voltage ramps from-100 mV to +100 mV in 500 ms (1 V·s^−1^) applied every 3 seconds from a holding potential 0 mV. The liquid-junction potential was calculated to be +4.9 mV using Clampex 10.4 software; data were not corrected for this offset. The recordings were performed at room temperature. A system for rapid superfusion was used, as described in Dittert et al.^91^, to wash the cells with extracellular bath solution containing: 160 mM NaCl, 2.5 nmM KCl, 1 mM CaCl_2_, 2 mM MgCl_2_, 10 mM HEPES, 10 mM glucose; pH adjusted to 7.3 with NaOH, 310 mosmol·l^−1^. The glass pipettes were filled with intracellular solution containing: 145 mM CsCl, 3 mM CaCl_2_, 2 mM MgATP, 10 mM HEPES, 5 mM EGTA; pH adjusted to 7.3 with CsOH, 300 mosmol·l^-1^. (-)-englerin A (EA; Phytolab) was dissolved in DMSO and stored as 1 mM aliquots at-80 °C. Before adding to cells, EA was diluted in the extracellular bath solution to a concentration of 30 nM (final DMSO concentration ∼0.003%). Reagents were purchased from Merck Life Science unless stated otherwise.

Electrophysiology data were analysed using Clampfit 10 and 11 (Molecular Devices, San Jose, CA, USA), SigmaPlot 10 (Systat Software Inc., San Jose, CA, USA) and OriginPro 2021 (OriginLab Corporation, Northampton, MA, USA). Voltage-dependent gating parameters were estimated from steady-state conductance-voltage (*G*/*V*) relationships obtained at the end of 100 ms voltage steps by fitting the conductance *G* = I/(*V*−*V*_rev_) as a function of the test potential *V* to the Boltzmann equation: *G* = ((*G*_max_ − *G*_min_)/(1 + exp[−*zF*(*V*−*V*_50_)/*RT*])) + *G*_min_, where *z* is the apparent number of gating charges involved in channel opening (in elementary charge units: *e*_o_ = 1.6 × 10^-19^ C), *V*_50_ is the half-activation voltage, *G*_min_ and *G*_max_ are the minimum and maximum whole-cell conductance, *V*_rev_ is the reversal potential, and *F*, *R*, and *T* have their usual thermodynamic meaning.

Throughout, average data are presented as means ± standard error of the mean (SEM), or as a median, range, and interquartile range as appropriate. Statistical significance was calculated using Student’s *t*-test, Mann-Whitney rank-sum, or one-way analysis of variance followed by the non-parametric Dunn’s test, as appropriate. Differences were considered significant at *P* < 0.05.

### Intracellular calcium recordings

HEK293 cells (ATCC) were cultured in Dulbecco’s Modified Eagle Medium (DMEM; Gibco^TM^) supplemented with 9% fetal bovine serum (Merck) and penicillin-streptomycin (Gibco^TM^). Cells were grown at 37 °C and 5 % CO_2_ and were tested to be negative for mycoplasma. For [Ca^2+^]_i_ recordings, cells were plated onto 6-well plates at 1 × 10^6^ cells/well and grown overnight. The following day, cells were transfected with 2 µg plasmid DNA and 6 µL JetPrime reagent (Polyplus) per well, according to manufacturer’s instructions. Control cells were transfected with 2 µg of empty vector (pcDNA4/TO). After 4 hours, transfection mixture was removed and replaced with fresh growth media. Cells were incubated overnight. 24 hours after transfection, cells were replated into poly-D-lysine (PDL; VWR) coated 96-well black, clear bottom plates (Nunc) at 50,000 cells/100 µl per well and allowed to adhere overnight. Cells were used for [Ca^2+^]_i_ recordings 48 hours after transfection.

For [Ca^2+^]_i_ recordings, medium was removed and cells incubated with SBS containing 2 µM Fura-2 AM (Molecular Probes) and 0.01% pluronic-F127 (Merck) for 60 minutes at 37 °C. SBS contained NaCl 135 mM, KCl 5 mM, HEPES 10 mM, glucose 8 mM, MgCl_2_ 1.2 mM, CaCl_2_ 1.5 mM, titrated to pH 7.4 with NaOH. Following incubation, Fura-2 AM was removed and 100 µl SBS was added to cells. Cells were incubated in SBS for 30 minutes at RT to allow de-esterification of Fura-2 AM. After incubation, SBS was removed and replaced with recording buffer (SBS containing vehicle to match compound buffer) prior to recording. [Ca^2+^]_i_ recordings were carried out using a FlexStation3 (Molecular Devices). Fura-2 fluorescence was measured with excitation wavelengths of 340 and 380 nm, and an emission wavelength of 510 nm. Measurements were taken at 5 s intervals for 5 minutes. At 60 s, compounds were automatically dispensed from a compound plate. For EA and AM237, compounds and recording buffer contained 0.01% pluronic F127.

Raw data were converted into a ratio of response at 340 nm to 380 nm. Responses were calculated at 250-300 s (unless stated otherwise), and the baseline at 0-55 s (before compound addition) was subtracted from these values. EC_50_ curves were fitted as variable slope (4 parameters). GraphPad Prism 10 and MS Excel were used for data processing and visualisation.

### Surface biotinylation

For surface biotinylation experiments, HEK293 cells were plated at 0.6 x 10^6^ cells/well into PDL-coated 6-well plates and allowed to adhere overnight. The next day, cells were transfected as described for [Ca^2+^]_i_ recordings. Transfections mixture was removed after 4 hours and replaced with fresh growth medium.

Surface biotinylation was carried out 48 hours after transfection. Medium was removed from cell and cells were washed 3× with ice-cold D-PBS (Merck) containing 1 mM CaCl_2_ to maintain cell adhesion. Cells were then incubated with D-PBS containing 1 mM CaCl_2_ and 0.3 mg/ml EZ-Link^TM^ Sulfo-NHS-LC-Biotin (ThermoFisher Scientific) for 30 minutes at 4 °C on a rocker. After this, excess biotin was quenched with D-PBS containing 1 mM CaCl_2_ and 100 mM glycine. Cells were lysed in NP-40 lysis buffer (ThermoFisher Scientific) and centrifuged to remove insoluble material. Protein in supernatants was quantified using the BCA Rapid Gold Assay (ThermoFisher Scientific), and 30 µg was removed for input samples. Input samples were prepared for SDS-PAGE with 4× Laemmli sample buffer (BioRad; supplemented with 10% β-mercaptoethanol) and heating at 95 °C for 10 minutes.

For streptavidin pulldowns, 500 µg of clarified lysate was incubated overnight at 4 °C with Magnetic Streptavidin Beads (Pierce^TM^; 25 µl slurry/sample) with rotation. Following washes with NP-40 lysis buffer, proteins were eluted off magnetic beads in 4× Laemmli sample buffer (supplemented with 10% β-mercaptoethanol) and 2 mM biotin at 95 °C for 10 minutes. Inputs and elutions were separated by SDS-PAGE on 7.5% Mini-PROTEAN® TGX™ Precast Gels (BioRad) and transferred to PVDF membrane. Membranes were blocked with 5% milk in PBS-Tween (PBS-T) before incubation with primary antibody against GFP (mouse; 1:5000, abcam,ab1218), overnight at 4 °C. For normalisation, surface blots were probed with a primary antibody against CD71 (rabbit; 1:5000, Cell Signalling Technology, #13113) and inputs blots with primary antibody against β-actin (mouse1:10,000; ThermoFisher # MA1-140). Primary antibodies were detected using anti-mouse HRP or anti-rabbit HRP secondary antibodies (1:5000, ThermoFisher Scientific), with incubation at RT for 60 min prior to PBS-T washes and detection using Pierce^TM^ ECL Western Blotting Substrate and iBright FL1500 imaging system (ThermoFisher Scientific).

## Supporting information

Extended Data Movie 1. TRPC5 structural dynamics.

## Supplementary Information

The following details can be found in the **Supplementary Information**: **Extended Data Movie 1**, highlighting the structural dynamics represented by our TRPC5:EA structures; **Supplementary Figure 1**, containing entire western blots for surface biotinylation experiments.

## Acknowledgements

Work at the University of Leeds was supported by Biotechnology and Biological Sciences Research Council (BBSRC) grants to RSB/SPM/DJB (BB/P020208/1; BB/Z514925/1) and a British Heart Foundation 4-year PhD studentship to KLRH. Large-scale tissue culture was performed in the University of Leeds Protein Production Facility, funded by the University of Leeds and the Royal Society (WL150028). We thank the Astbury Biostructure Laboratory, funded by the University of Leeds and Wellcome (108466/Z/15/Z; 221524/Z/20/Z; 218785/Z/19/Z), for support of electron microscopy work. Work at the Institute of Physiology, Prague was supported by the Grant Agency of the Czech Republic (GACR 24-10147S). Professor Eric Gouaux (Vollum Institute) is kindly acknowledged for providing the BacMam vector. We thank Dr Charlotte A. Scarff (University of Leeds) for providing critical comments on our manuscript.

## Author Contributions

SAP performed protein production and cryo-EM studies, including grid preparation, data collection and processing, model building and structural analysis. AP performed mutagenesis and patch-clamp electrophysiology. CCB performed mutagenesis, surface biotinylation and intracellular calcium measurements. KLRH contributed to model building. SAP, AP, CCB, KLRH, VV, SPM and RSB analysed and interpreted data. SAP, AP, VV, CCB, KLRH and RSB prepared figures. DJB, VV, SPM and RSB supervised experiments. SAP, KLRH, DJB, SPM, VV and RSB secured funding. DJB, SPM, VV and RSB conceptualised the project. RSB led the project. SAP, VV and RSB drafted the paper with input from CCB and AP. All authors commented on the paper draft.

## Competing Interest

RSB and DJB are scientific co-founders and partners of the pharmaceutical start-up company CalTIC GmbH. RSB, DJB and SAP have received funding from CalTIC GmbH. DJB is an inventor on the following patent applications: (1) PCT/GB2018/050369. TRPC ion channel inhibitors for use in therapy. Filing date: 9th February 2018; (2) 62/529,063. Englerin derivatives for the treatment of cancer. Filing date: 6th July 2017. The other authors declare no competing interest.

## Availability of data and materials

Cryo-EM models and maps are available via the PDB and EMDB, respectively (**TRPC5^PA^:** 9RRF / EMD-54186, **TRPC5:EA^PA^-S1:** 9RRM / EMD-54187, **TRPC5:EA^PA^-S2:** 9RRN / EMD-54188, **TRPC5:EA_4:4_-S1:** 9RRU / EMD-54204, **TRPC5:EA_4:2_-S1:** 9RRQ / EMD-54193, **TRPC5:EA_mix_-S1:** 9RRO / EMD-54189, **TRPC5:EA_4:2_-S2:** 9RVV / EMD-54291, **TRPC5:EA_mix1_-S2:** 9RSH / EMD-54219, **TRPC5:EA_mix2_-S2:** 9RSG / EMD-54218). Details of cryoEM structures are provided in **Extended Data Table 1**. Example data for intracellular calcium recordings, patch-clamp electrophysiology and surface biotinylation experiments are displayed in **Figure 2**, **Figure 4**, **Figure 5** and **Extended Data Figures 4-6**. Full western blots for surface biotinylation experiments are provided in **Supporting Figure 1**. Other data and materials are available from the corresponding authors upon reasonable request.

## Extended Data Figures, Movies and Tables

**Extended Data Figure 1.**
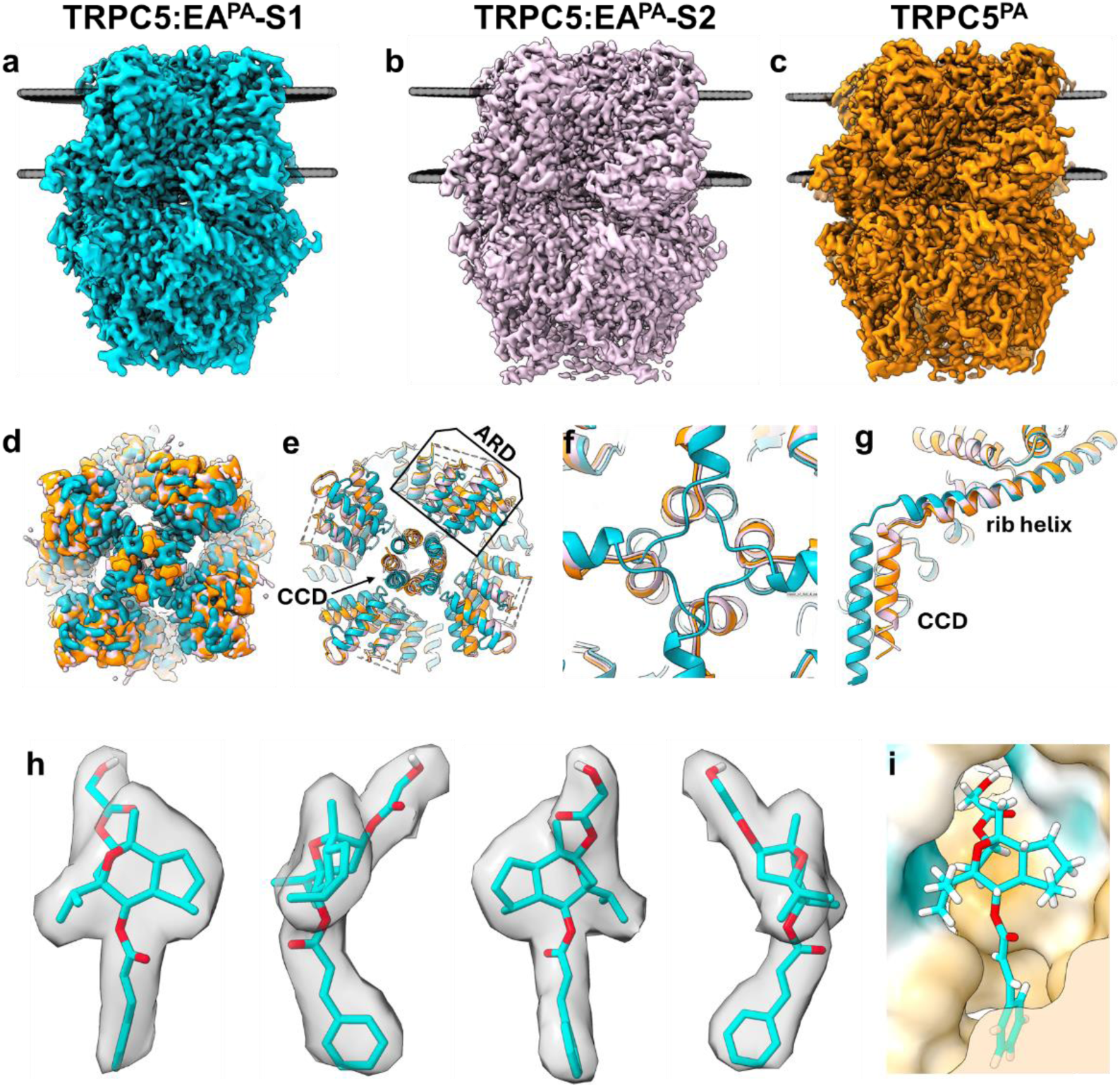
Cryo-EM structures reveal multiple ARD states and the EA binding site of the human TRPC5 channel. a-c, cryo-EM maps of TRPC5:EA^PA^-S1, TRPC5:EA^PA^-S2 and TRPC5^PA^. **d,e,** superimposition of cryo-EM maps and models of TRPC5:EA^PA^-S1 (blue) and TRPC5:EA^PA^-S2 (orange) (bottom views) showing the main differences between ARD states. **f,** Close-up of the superimposed lower gates of TRPC5:EA^PA^-S1 (blue) and TRPC5:EA^PA^-S2 (orange) (bottom views). **g,** Side view of the superimposed CCD and rib helix of TRPC5:EA^PA^-S1 (blue) and TRPC5:EA^PA^-S2 (orange). **h,** Multiple viewing angles of EA fitted in the EM density of TRPC5:EA^PA^-S1. **i,** Hydrophobic environment of the EA binding coloured by lipophilicity, with dark cyan being the most hydrophilic and goldenrod being the most lipophilic surface.

**Extended Data Figure 2.**
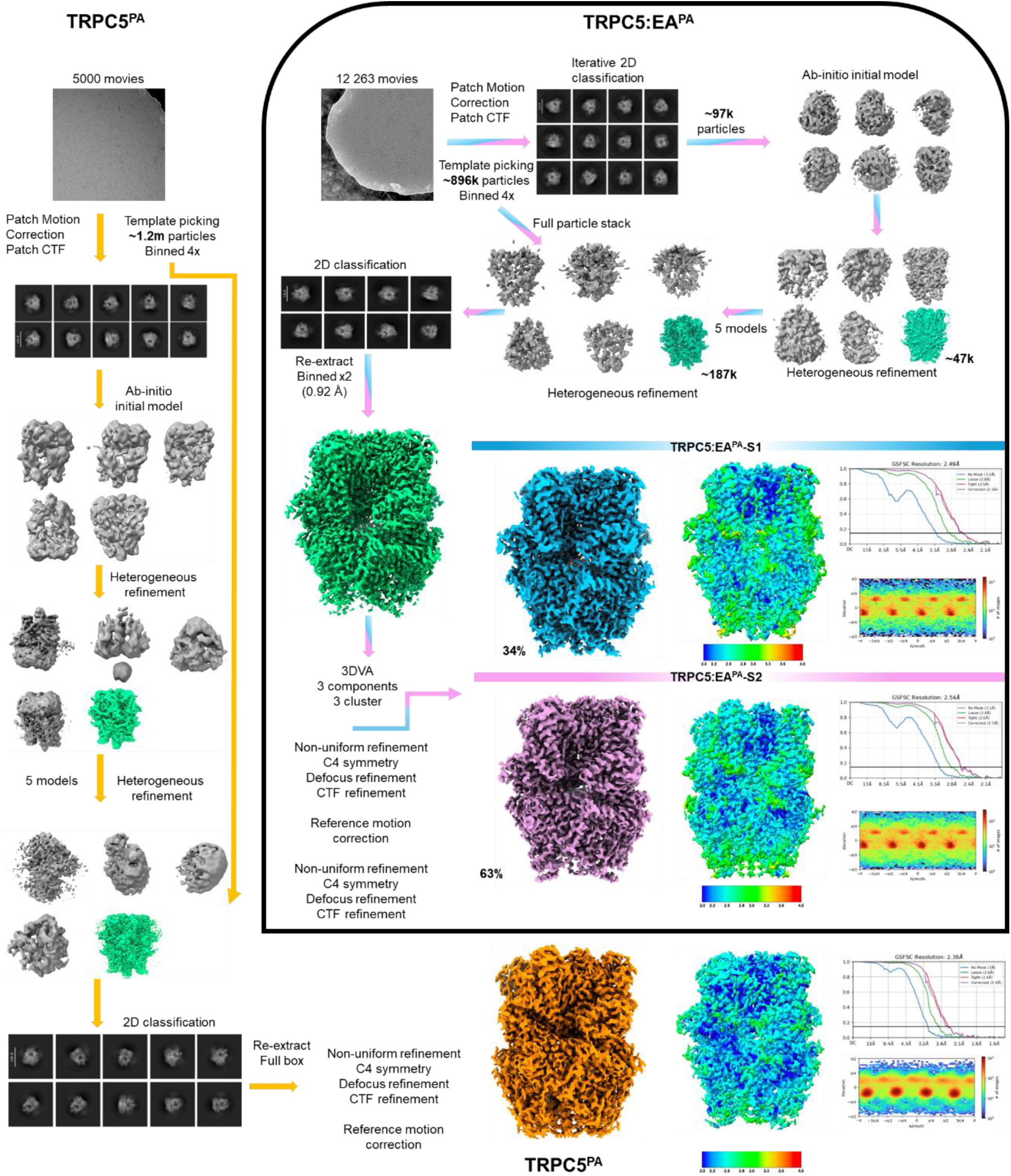
Cryo-EM data processing workflow and map resolution of TRPC5:EA^PA^-S1, TRPC5:EA^PA^-S2 and TRPC5^PA^.

**Extended Data Figure 3.**
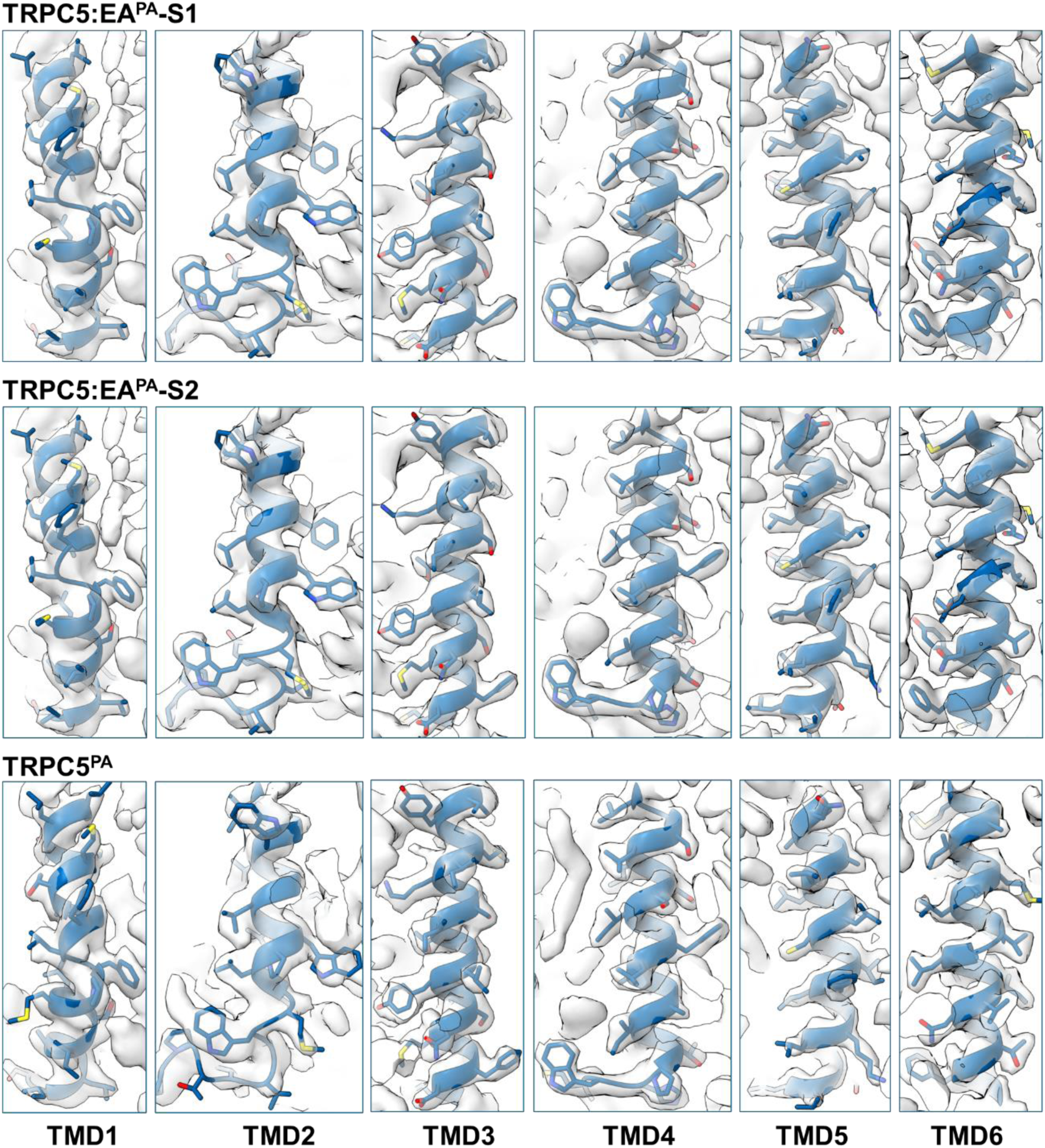
**Data quality of TRPC5:EA^PA^-S1, TRPC5:EA^PA^-S2 and TRPC5^PA^ illustrated by the fit of the six transmembrane domains (blue) to the EM maps (grey).**

**Extended Data Figure 4.**
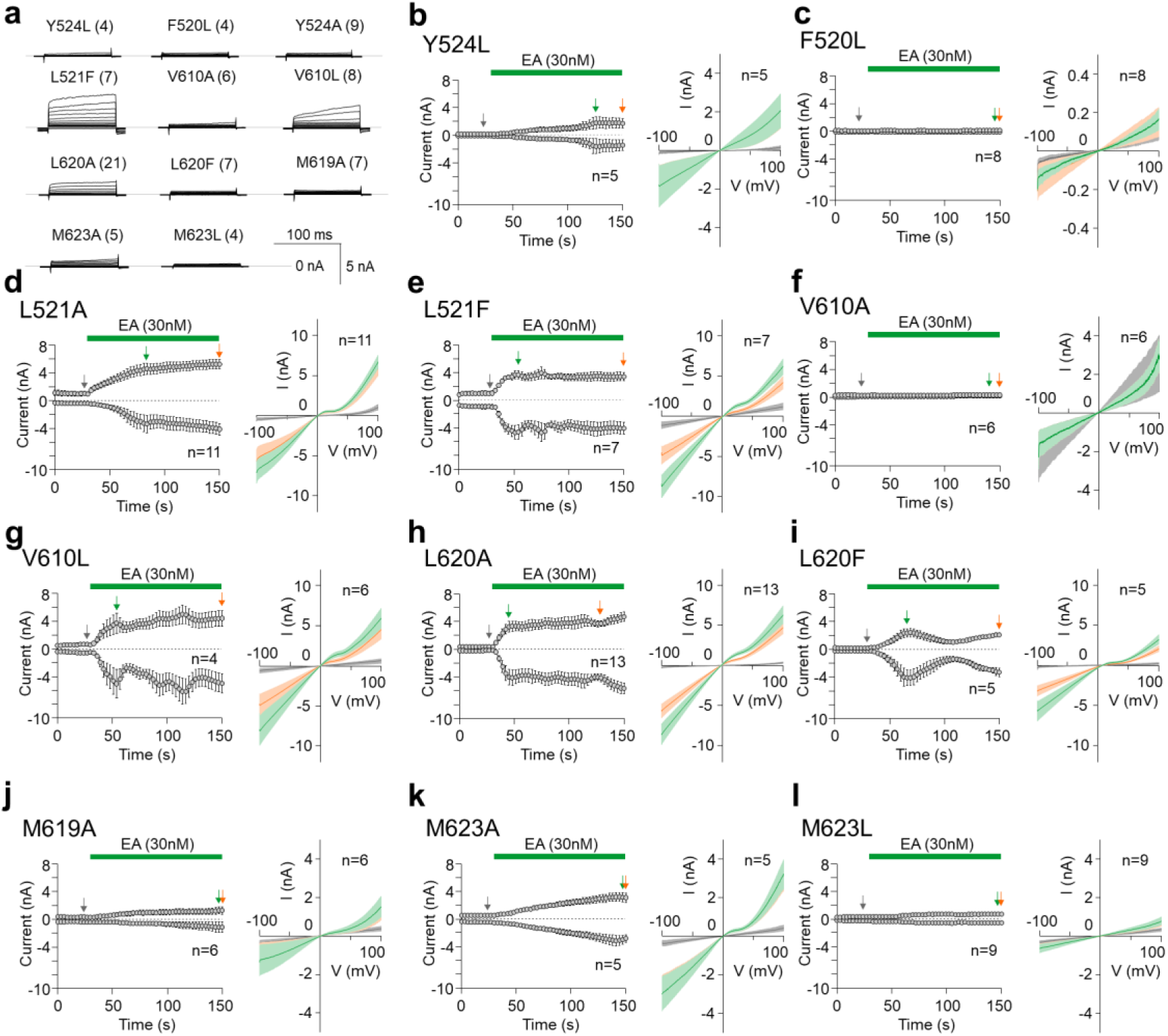
Functional analysis of TRPC5 variants. a, Mean current trace in response to 100-ms voltage steps from-80 to +200 mV (20 mV increment) recorded from HEK293T cells expressing indicated TRPC5 constructs. The currents were recorded ∼1 min after whole-cell formation in extracellular control solution. Numbers of cells (n) are indicated in parentheses. **b-l,** Time courses of average whole-cell currents elicited by 30 nM EA in HEK293T cells expressing indicated constructs of TRPC5. A ramp pulse from-100 mV to +100 mV was periodically applied from a holding potential of 0 mV every 3 seconds for 500 ms. Amplitudes were measured at-100 mV and +100 mV and the mean ± SEM was plotted as a function of time. Numbers of cells (n) are indicated. Right panels for each construct: mean current-voltage relations (coloured curves, ± SEM as lighter-coloured envelopes) are plotted for the currents measured at times indicated in the left panel by vertical arrows (grey at baseline, green at peak, and orange after 2-min exposure to EA).

**Extended Data Figure 5.**
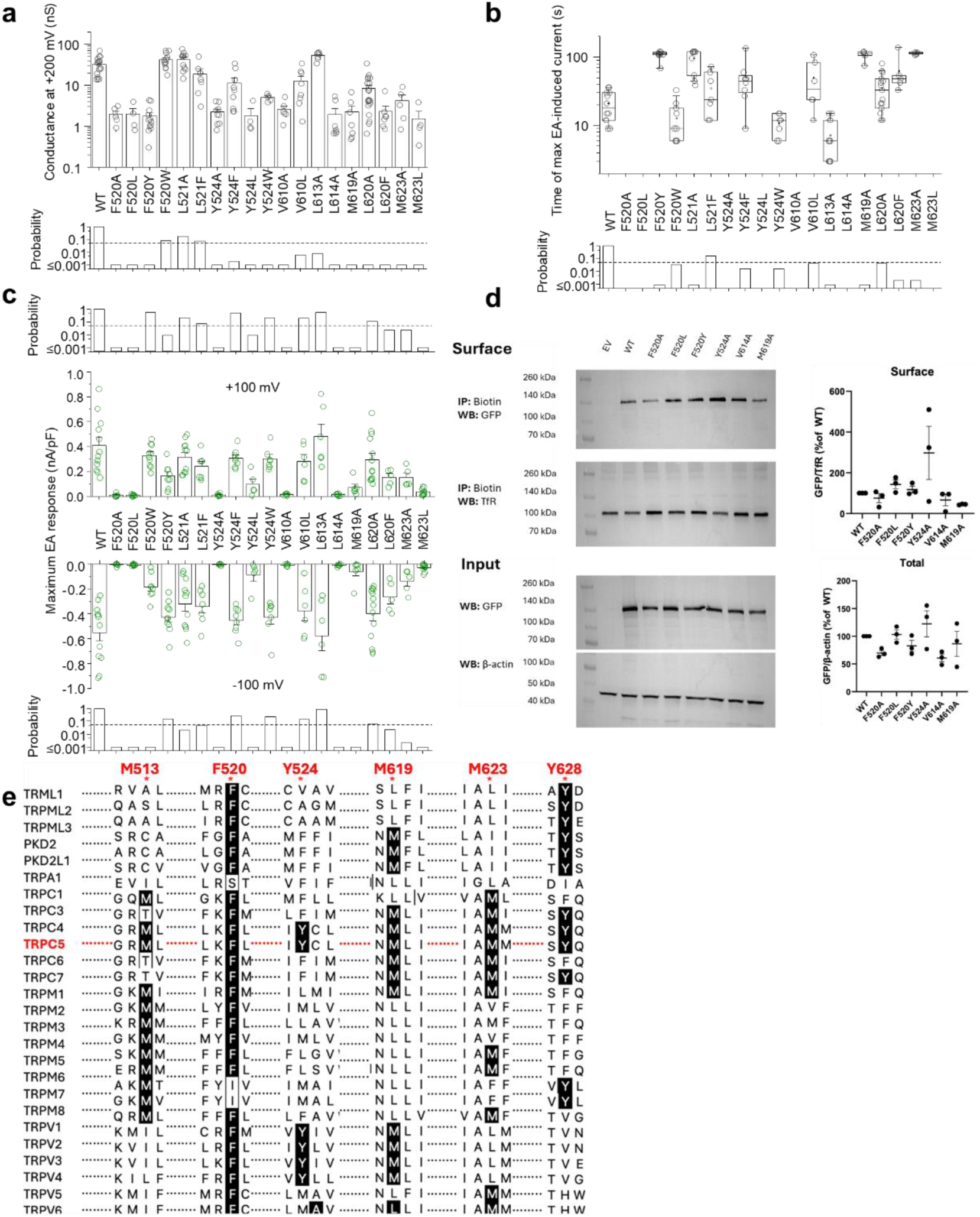
Effects of mutagenesis of residues surrounding the EA binding site of TRPC5. a, Summary of effects of TRPC5 mutagenesis on voltage-induced TRPC5 activation, showing average currents at +200 mV obtained from the protocol shown in **Figure 4d**. Below, bar graph representing the probabilities obtained from the Student’s two-sided unpaired t-tests, performed to determine if there was a significant difference between the responses of the wild-type and the individual constructs. The level of statistical significance (P < 0.05) is indicated with a dashed horizontal line. **b,** Box plot of values of times to maximal responses mediated by the indicated constructs within 2 min of 30 nM EA application, measured at +100 mV. For each box, the centre line is the median value, square is the mean, the edges of the boxes are the 25^th^ and 75^th^ percentiles, and the lines extending from the boxes are the 5^th^ and 95^th^ percentiles (n ≥ 5). **c,** Bar graphs representing summarised current densities of the maximal responses measured within 2 min of 30 nM EA application at +100 mV and-100 mV for indicated TRPC5 constructs. Above and below, bar graphs representing the probabilities obtained from the Student’s two-sided unpaired t-tests, performed to determine if there was a significant difference between the EA responses of the wild-type and the mutated constructs. The level of statistical significance (P < 0.05) is indicated with a dashed horizontal line. **d,** Surface biotinylation experiments of TRPC5-SYFP2 variants expressed in HEK293 cells. Biotinylated surface proteins (upper blots) were pulled down using magnetic streptavidin beads and probed with antibodies against GFP (TRPC5-SYFP2) and Transferrin Receptor (TfR; loading control). Input samples were blotted with antibody against GFP (TRPC5-SYFP2) and β-actin (loading control) to confirm expression (lower blots). Blots were analysed using densitometry analysis and expression compared to control (WT TRPC5-SYFP2). Data are displayed as mean ± SEM. Data were analysed using one-way ANOVA with Dunnett’s multiple comparison test to compare mutants to WT (control) **e,** Conservation analysis of key residues involved in aromatic interactions changes induced by EA binding in TRPC5. hTRPC5 residues were aligned with the sequences of other human TRP members in the MEGA (v11.0.6)^92^.

**Extended Data Figure 6.**
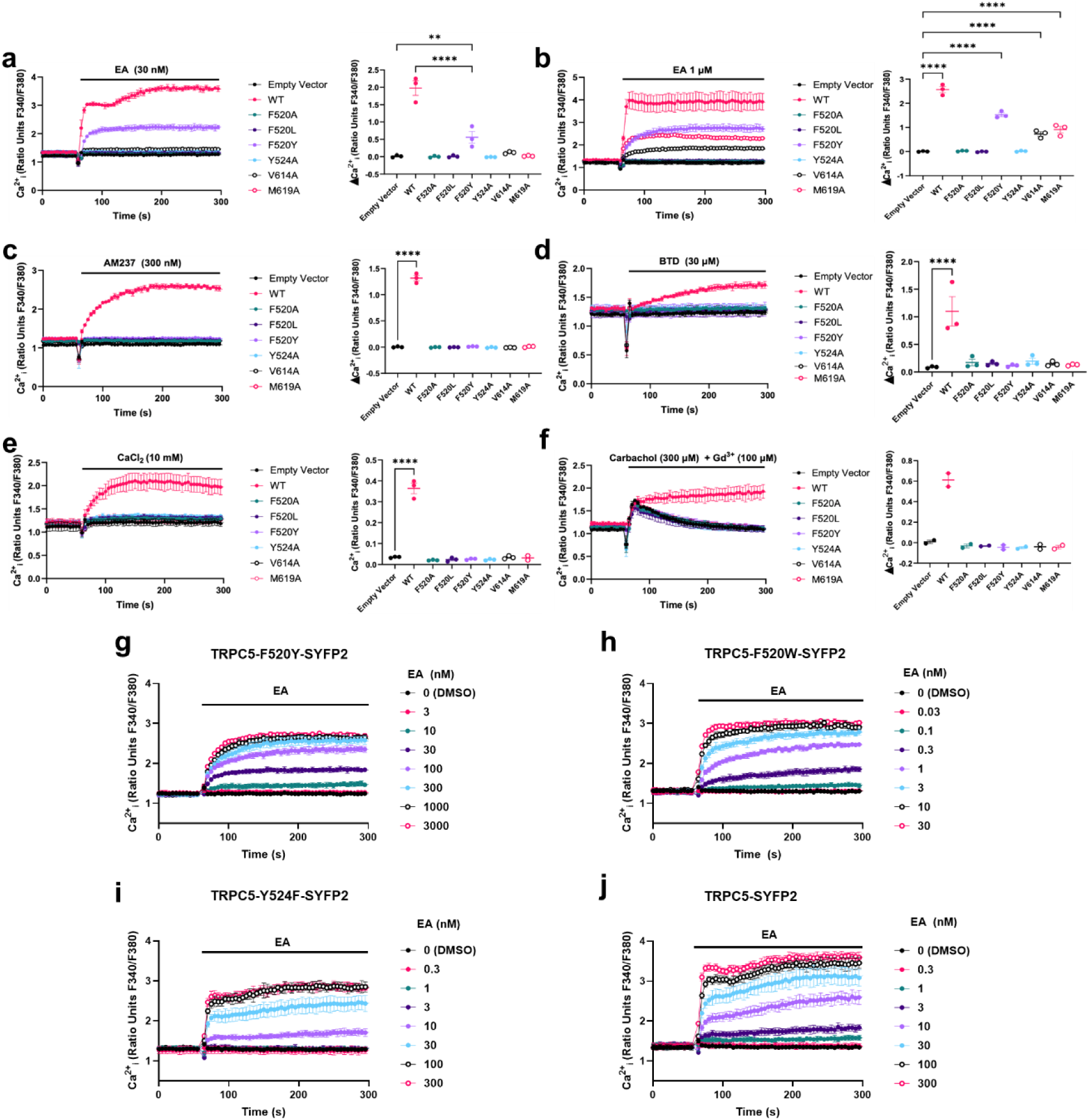
Intracellular Ca^2+^ recordings of TRPC5-SYFP2 variants. a-f, Left, Representative traces from one 96-well plate (N = 6 technical repeats) each, showing increase in [Ca^2+^]_i_ of HEK293 cells expressing indicated TRPC5-SYFP2 variants, in response to different TRPC5 activators: 30 nM EA (a), 1 µM EA (b), 300 nM AM237 (c), 30 µM BTD (d), 10 mM extracellular CaCl_2_ (e) or a combination of 300 µM carbachol and 100 µM GdCl_3_ (f). Data are shown as mean ± SD. Right, mean responses (± SEM, n = 3 independent experiments) of experiments shown on the left of each panel, calculated by subtracting the basal levels (at 0-55 s) from the activator-induced responses (at 250-300 s). Data were analysed using one-way ANOVA with Dunnett’s multiple comparison test to compare mutants to WT (control), and Šídák’s multiple comparisons test (for a only) **g-j,** Representative traces from one 96-well plate (N = 6 technical repeats) each, showing concentration-dependent increases in [Ca^2+^]_i_ in response to indicated concentrations of EA in HEK293 cells expressing TRPC5_F520Y_-SYFP2 (g), TRPC5_F520W_-SYFP2 (h), TRPC5_Y524F_-SYFP2 (i), and wild-type TRPC5-SYFP2 (j). Data are shown as mean ± SD. Corresponding concentration-response curves for (g-j) from 3 independent experiments (n = 3) are shown in **Figure 2l**.

**Extended Data Figure 7.**
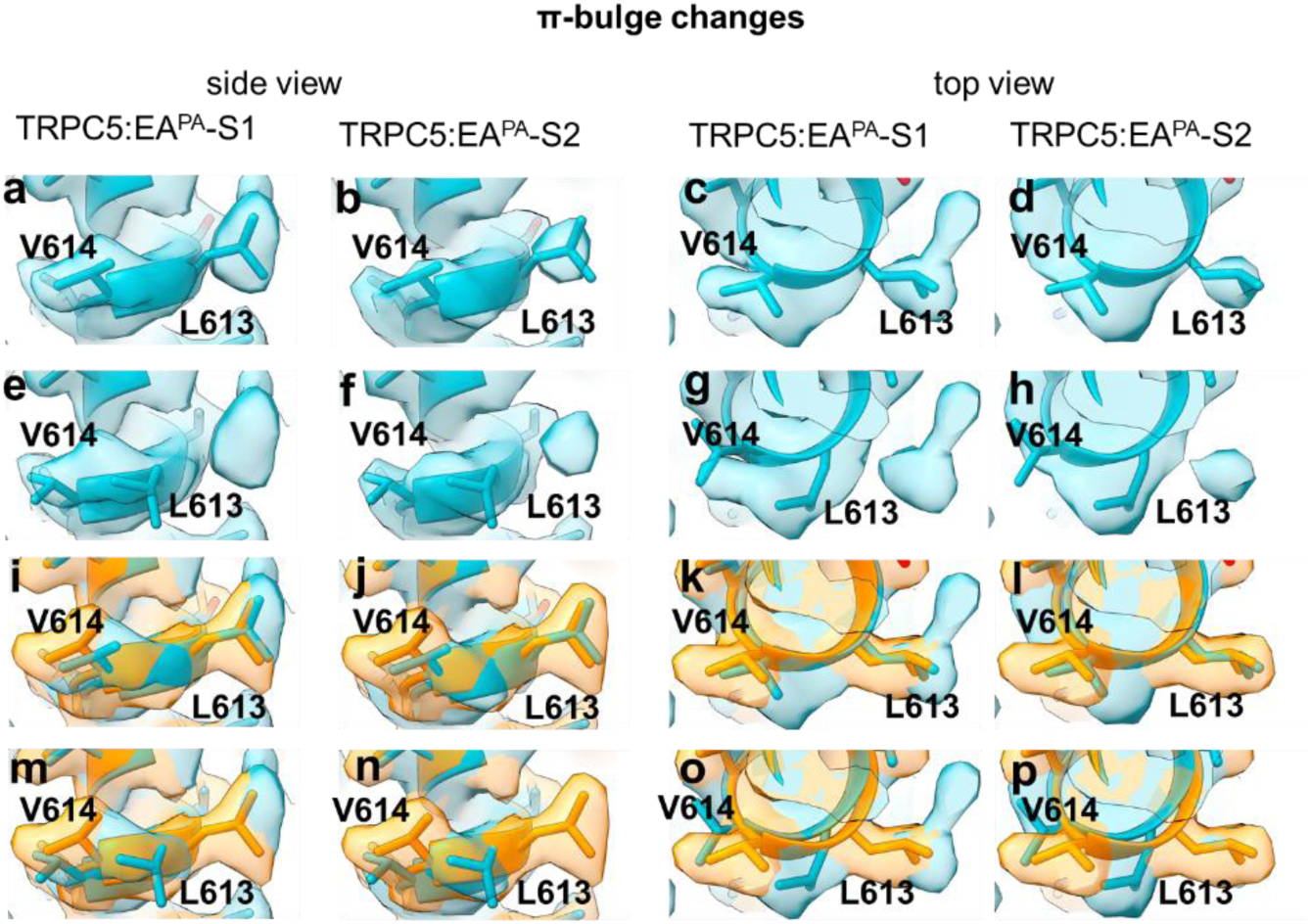
EA binding induces flexibility of the TRPC5 π-bulge. a-h, Ambiguous density around the TRPC5 π-bulge allows fitting of different rotameric position for L613 and V614 into the EM maps of TRPC5:EA^PA^-S1 (a,c,e,g) and TRPC5:EA^PA^-S2 (b,d,f,h), shown from the side (a,b,e,f) and top (c,d,g,h). **i-p,** Overlays of the maps and models of the TRPC5 π-bulge in TRPC5:EA^PA^-S1 and TRPC5:EA^PA^-S2 and the map and model of TRPC5^PA^ (orange), illustrating the differences in map certainty and suggesting EA-induced flexibility of this region.

**Extended Data Figure 8.**
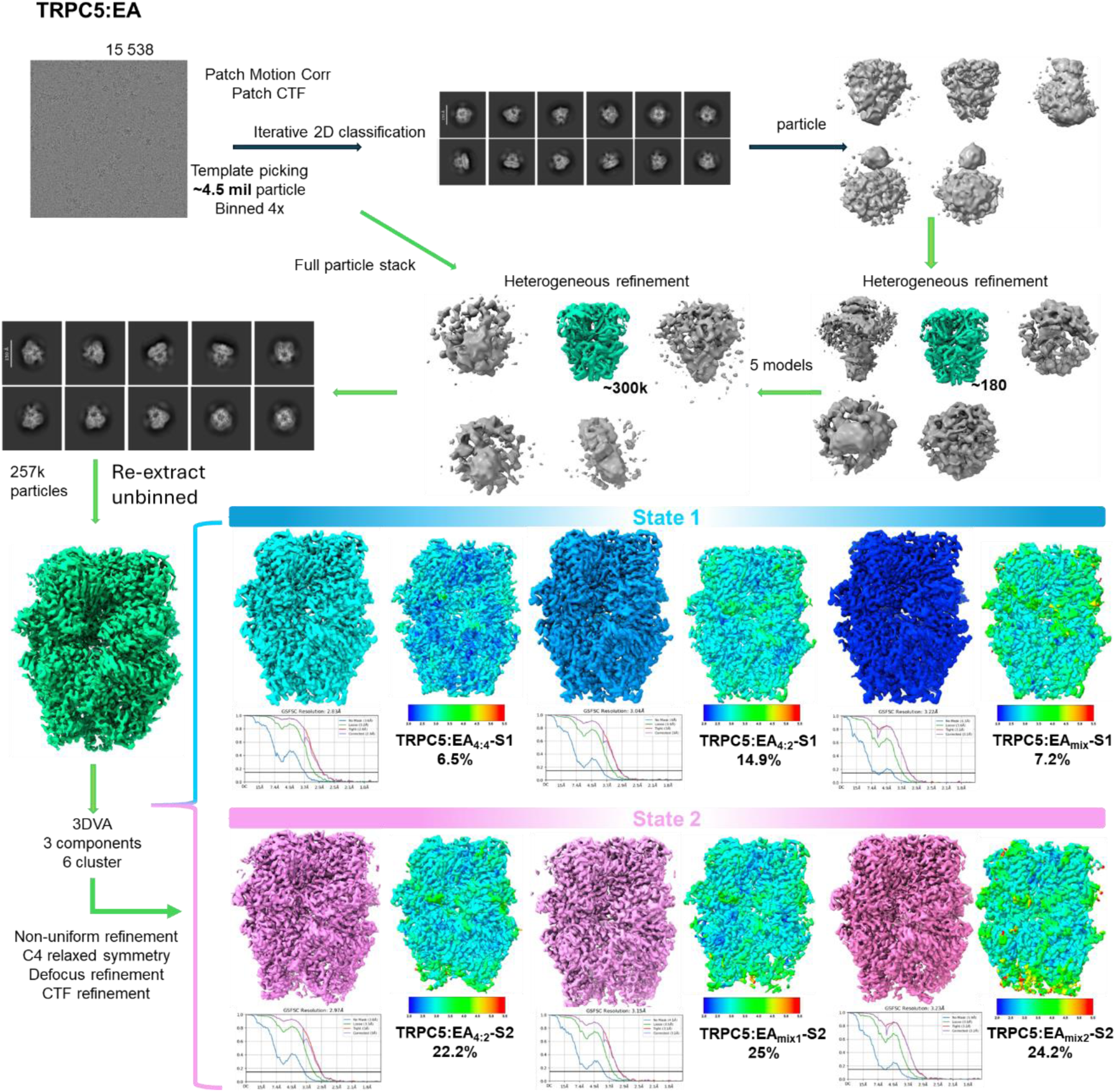
Cryo-EM data processing workflow and map resolution of TRPC5:EA structures.

**Extended Data Figure 9.**
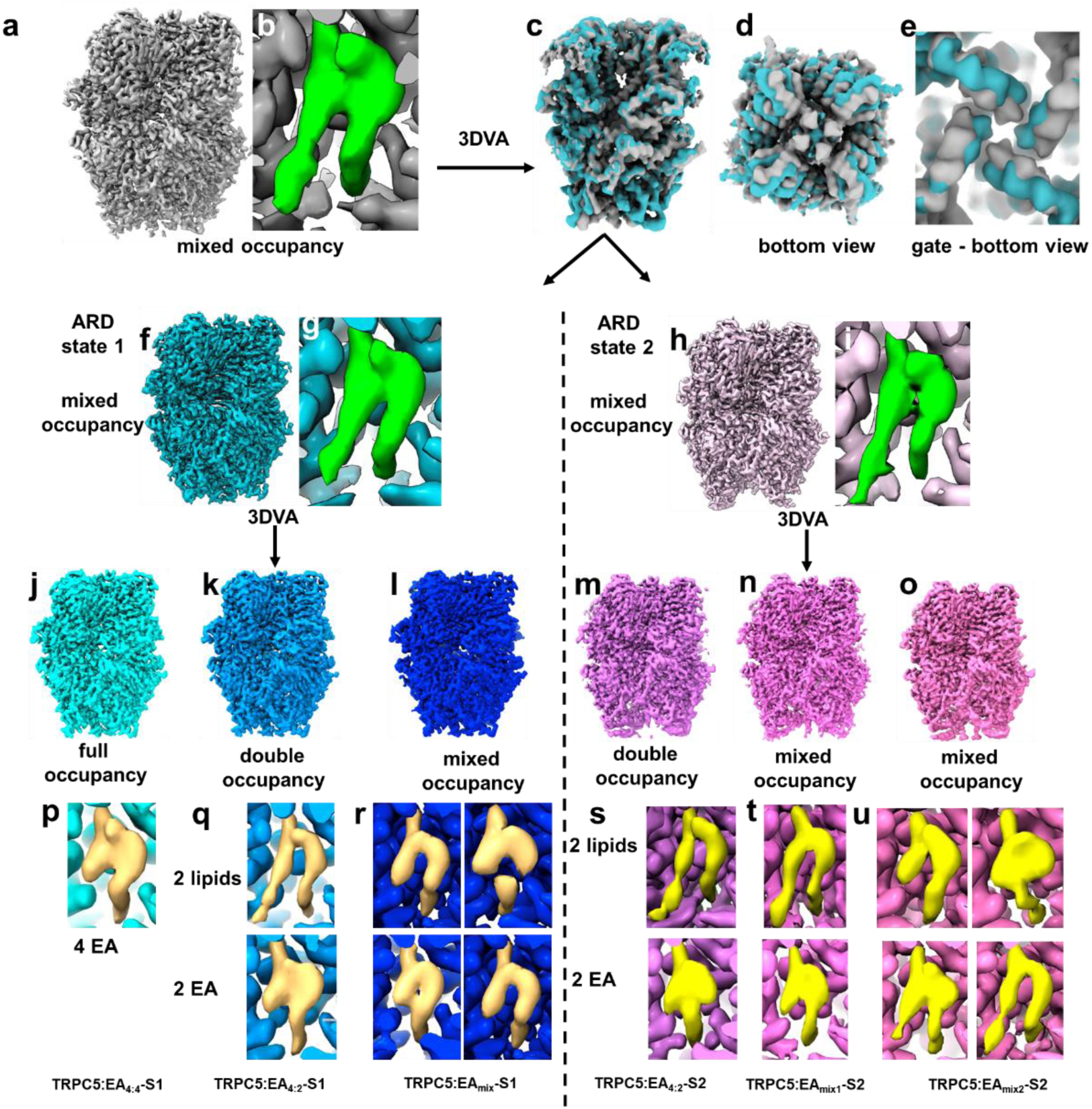
Structures of TRPC5:EA reveal multiple EA/lipid binding stoichiometries. a,b, Initial cryo-EM map of TRPC5:EA (a), with a close-up of the EA binding site showing ambiguous density (green; b)**. c-e,** 3DVA analysis and separation of two ARD states of TRPC5 illustrated with a side view (c), bottom view (d) and a close-up on the lower gate (e). **f-i,** Cryo-EM maps of ARD state 1 (f) and ARD state 2 (h) and close-ups on the respective EA binding sites (g,i) showing ambiguous density (green). **j-l,** ARD state 1 cryo-EM maps with different EA/lipid binding stoichiometries (4:4 in cyan; 4:2 in light blue; mixed in dark blue). **m-o,** ARD state 2 cryo-EM maps with different EA/lipid binding stoichiometries (4:2 in magenta; mixed1 in light magenta; mixed2 in pink). **p-r,** Close-ups of EA binding sites of ARD state 1 maps showing different non protein densities (cream). **s-u,** Close-ups of EA binding sites of ARD state 2 maps showing different non protein densities (yellow).

**Extended Data Figure 10.**
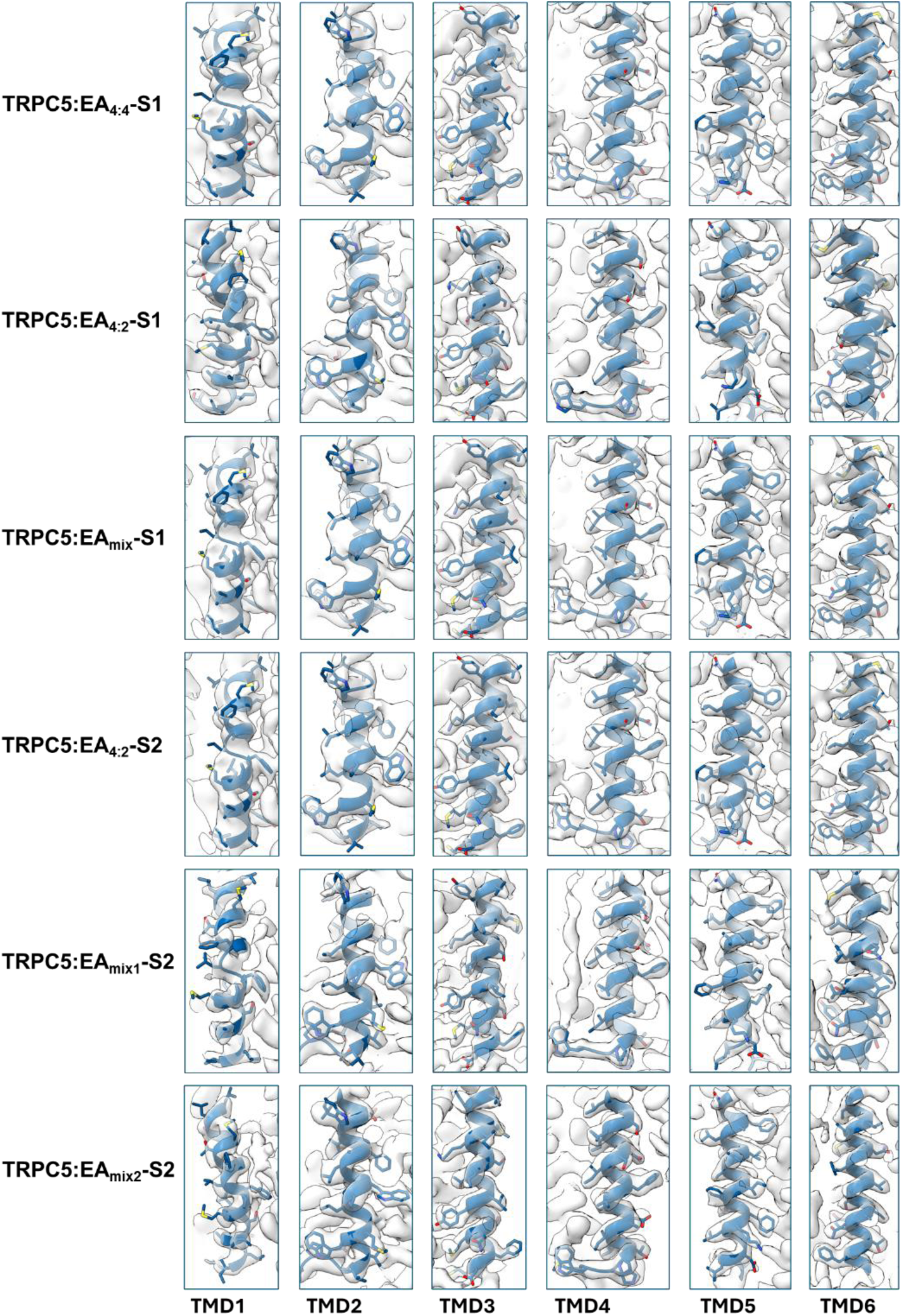
Data quality of TRPC5:EA structures illustrated by the fit of the six transmembrane domains (blue) in the EM maps (grey).

**Extended Data Figure 11.**
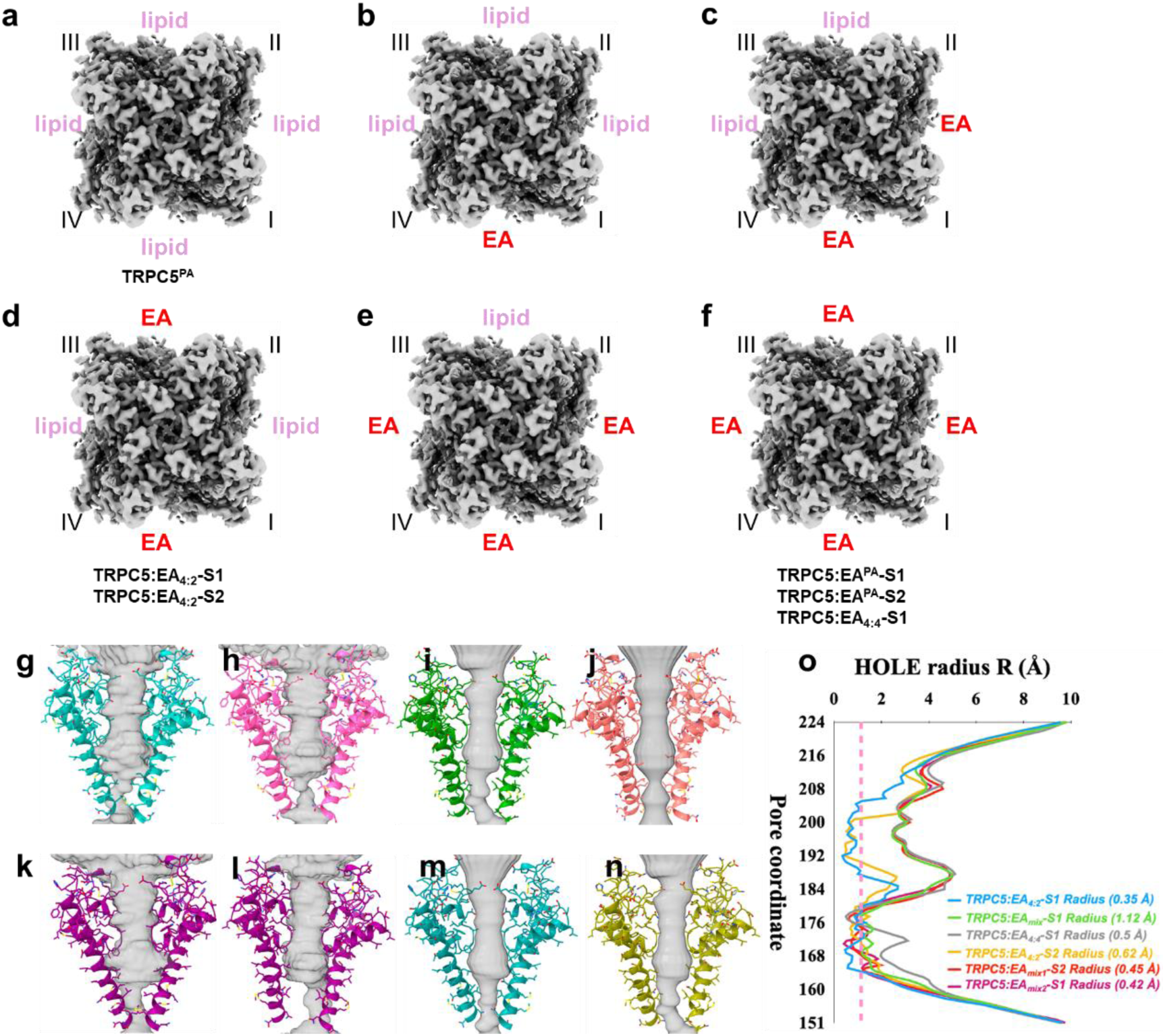
Stoichiometry and pore analysis of TRPC5:EA structures. a-f, Possible stoichiometries of EA:lipid in TRPC5. Structures of three possible arrangements were determined unambiguously: TRPC5^PA^ (a); TRPC5:EA**_4:2_**-S1 and TRPC5:EA**_4:2_**-S2 (d); TRPC5:EA^PA^-S1, TRPC5:EA^PA^-S2 and TRPC5:EA**_4:4_**-S1 (f). Three additional structures (TRPC5:EA**_mix_**-S1, TRPC5:EA**_mix1_**-S2 and TRPC5:EA**_mix2_**-S2 represent mixed populations of arrangements shown in panels (b-e) and could not be separated further. **g-h,** Pore shape analysis of TRPC5:EA**_4:2_**-S1, with projections showing TRPC5 subunits I and III (cyan; g) and TRPC5 subunits II and IV (pink; h). **i,j,** Pore shape analysis of TRPC5:EA**_mix_**-S1 (i), and TRPC5:EA**_4:4_**-S1 (j). **k,l,** Pore shape analysis of TRPC5:EA**_4:2_**-S2, with projections showing TRPC5 subunits I and III (k) and TRPC5 subunits II and IV (l). **m,n,** Pore shape analysis of TRPC5:EA**_mix1_**-S2 (m) and TRPC5:EA**_mix2_**-S2 (n). **o,** Plot of the pore radii of TRPC5 structures against pore coordinates down the channels. The pink dashed line indicates where the pore becomes too narrow for water to pass through (1.2 Å). Pore radius of TRPC5:EA structures. Pore shape analysis and calculation of pore radii was performed using PoreAnalyzer. **Extended Data Movie 1. TRPC5 structural dynamics.** Our TRPC5 data set was analysed using 3DVA and the output was visualised using ‘simple mode’ in CryoSPARC. The resulting movie shows transitions between the ARD states of TRPC5 and the symmetry break in the channel pore.

---

**Extended Data Table 1.** Cryo-EM data collection, refinement and validation statistics.

| **PDB**<br>**EMDB** | **TRPC5<sup>PA</sup>**<br>**9RRF**<br>**EMD-54186** | **TRPC5:EA<sup>PA</sup>-S1**<br>**9RRM**<br>**EMD-54187** | **TRPC5:EA<sup>PA</sup>-S2**<br>**9RRN**<br>**EMD-54188** | **TRPC5:EA<sub>4:4</sub>-S1**<br>**9RRU**<br>**EMD-54204** | **TRPC5:EA<sub>4:2</sub>-S1**<br>**9RRQ**<br>**EMD-54193** | **TRPC5:EA<sub>mix</sub>**<br>**9RRO**<br>**EMD-54189** | **TRPC5:EA<sub>4:2</sub>-S2**<br>**9RVV**<br>**EMD-54291** | **TRPC5:EA<sub>mix1</sub>-S2**<br>**9RSH**<br>**EMD-54219** | **TRPC5:EA<sub>mix2</sub>-S2**<br>**9RSG**<br>**EMD-54218** |
| --- | --- | --- | --- | --- | --- | --- | --- | --- | --- |
| **Data collection and processing** |  |  |  |  |  |  |  |  |  |
| Magnification | 165k | 270k | 270 | 165k | 165k | 165k | 165k | 165k | 165k |
| Voltage (kV) | 300 | 300 | 300 | 300 | 300 | 300 | 300 | 300 | 300 |
| Electron exposure<br>(e<sup>-</sup>/Å<sup>2</sup>) | 35.46 | 40.65 | 40.65 | 34.75 | 34.75 | 34.75 | 34.75 | 34.75 | 34.75 |
| Defocus range (μm) | -0.7 to -3.0 | -0.7 to -3.0 | -0.7 to -3.0 | -0.7 to -3.0 | -0.7 to -3.0 | -0.7 to -3.0 | -0.7 to -3.0 | -0.7 to -3.0 | -0.7 to -3.0 |
| Pixel size (Å) | 0.74 | 0.46 | 0.46 | 0.82 | 0.82 | 0.82 | 0.82 | 0.82 | 0.82 |
| Symmetry imposed | C4 | C4 | C4 | C4 | C4 relaxed | C1 | C1 | C1 | C1 |
| Final particle images<br>(no.) | 167,808 | 37,807 | 69,865 | 16,767 | 38,365 | 18,655 | 57,236 | 63,619 | 62,384 |
| Map resolution (Å) | 2.4 | 2.5 | 2.5 | 2.8 | 3.0 | 3.2 | 3.0 | 3.1 | 3.2 |
| FSC threshold | 0.143 | 0.143 | 0.143 | 0.143 | 0.143 | 0.143 | 0.143 | 0.143 | 0.143 |
| **Refinement** |  |  |  |  |  |  |  |  |  |
| Initial model | ModelAngelo | ModelAngelo | ModelAngelo | ModelAngelo | ModelAngelo | ModelAngelo | ModelAngelo | ModelAngelo | ModelAngelo |
| Map sharpening *B* factor<br>(Å<sup>2</sup>) | -77.7 | -59.6 | -71.4 | -57.0 | -48.5 | -36.9 | -57.2 | -60.2 | -56.4 |
| *Model composition* |  |  |  |  |  |  |  |  |  |
| Non-hydrogen atoms | 24182 | 23732 | 23240 | 23808 | 23615 | 22173 | 22623 | 208/08 | 20866 |
| Protein residues | 2800 | 2772 | 2716 | 2780 | 2749 | 27214 | 2630 | 2560 | 2563 |
| Ligands | 20 | 20 | 20 | 20 | 20 |  | 20 |  |  |
| *Bonds (RMSD)* |  |  |  |  |  |  |  |  |  |
| Length (Å) (# > 4σ) | 0.003(0) | 0.003(0) | 0.001(0) | 0.003(0) | 0.002(0) | 0.003 | 0.003(0) | 0.004(0) | 0.002(0) |
| Angles (°) (# > 4σ) | 0.417(0) | 0.461(0) | 0.343(0) | 0.426(0) | 0.417(0) | 0.658(2) | 0.498(0) | 0.632(3) | 0.537(0) |
| **Validation** |  |  |  |  |  |  |  |  |  |
| **MolProbity score** | 2.34 | 2.03 | 1.74 | 2.17 | 2.2 | 1.73 | 2.21 | 1.82 | 1.71 |
| Clash score | 17.72 | 16.87 | 17.36 | 18.84 | 23.24 | 9.92 | 23.37 | 8.23 | 7.79 |
| *Ramachandran plot (%)* |  |  |  |  |  |  |  |  |  |
| Outliers | 0.04 | 0.00 | 0.15 | 0.00 | 0.07 | 0.07 | 0.08 | 0.04 | 0.24 |
| Allowed | 4.32 | 3.07 | 1.49 | 2.26 | 3.13 | 3.23 | 3.27 | 4.84 | 3.84 |
| Favoured | 95.64 | 96.93 | 98.36 | 97.74 | 96.80 | 96.69 | 96.65 | 95.12 | 95.92 |
| Rotamer outliers (%) | 2.4 | 1.45 | 0.78 | 2.78 | 1.55 | 1.36 | 1.50 | 1.1 | 0.48 |
| *B factors* |  |  |  |  |  |  |  |  |  |
| Protein | 6.5/180.6/74.7 | 9.6/179.4/80.5 | 5.24/171.9/73.8 | 10.5/140.7/58.2 | 36.8/201.4/95.9 | 60.1/197.4/111.3 | 5.4/184.7/59.9 | 3.2/200.1/59.0 | 9.17/184.2/63.0 |
| Ligand | 18.9/177.7/59.2 | 25.2/170.5/70.1 | 17.5/153.8/58.1 | 24.3/143.7/56.7 | 58.5/206.2/90.7 |  | 12.6/168.4/37.3 |  |  |
| **Model vs. Data** |  |  |  |  |  |  |  |  |  |
| CC (mask) | 0.9 | 0.82 | 0.8 | 0.88 | 0.84 | 0.84 | 0.86 | 0.88 | 0.87 |
| CC (box) | 0.68 | 0.64 | 0.64 | 0.69 | 0.66 | 0.66 | 0.7 | 0.74 | 0.7 |
| CC (peaks) | 0.72 | 0.61 | 0.61 | 0.66 | 0.57 | 0.54 | 0.65 | 0.69 | 0.65 |
| CC (volume) | 0.87 | 0.79 | 0.77 | 0.85 | 0.83 | 0.83 | 0.82 | 0.85 | 0.85 |
| Mean CC for ligands | 0.7 | 0.67 | 0.7 | 0.67 | 0.66 |  | 0.71 |  |  |

**Extended Data Table 2.** Overview of TRPC5 residues that could not be modelled in TRPC5 structures.

| Structure | Subunits |  |  |  |
| --- | --- | --- | --- | --- |
|  | I | II | III | IV |
| TRPC5<sup>PA</sup> | 1-16, 120-133, 667-698, 763-765 |  |  |  |
| TRPC5:EA<sup>PA</sup>-S1 | 1-14, 120-134, 279-284, 666-699, 762-765 |  |  |  |
| TRPC5:EA<sup>PA</sup>-S2 | 1-16, 120-133, 276-284, 666-699, 753-765 |  |  |  |
| TRPC5:EA<sub>4:2</sub>-S1 | 1-15, 120-133, 276-284, 665-700, 763-765. | 1-15, 120-133, 276-284, 665-700, 763-765. | 1-15, 120-133, 276-284, 665-700, 763-765. | 1-15, 119-134, 276-284, 665-700, 762-765. |
| TRPC5:EA<sub>4:4</sub>-S1 | 1-14, 120-134, 279-284, 666-699, 765. |  |  |  |
| TRPC5:EA<sub>mix</sub>-S1 | 1-15, 119-134, 274-285, 665-702, 764-765. | 1-15, 119-134, 274-285, 665-703, 764-765. | 1-15, 119-134, 274-285, 665-702, 764-765. | 1-15, 119-134, 274-285, 630-632, 665-702, 764-765. |
| TRPC5:EA<sub>mix1</sub>-S2 | 1-15, 119-133, 275-284, 666-700, 733-765. |  |  |  |
| TRPC5:EA<sub>mix2</sub>-S2 | 1-15, 119-133, 277-284, 386-390, 666-701, 747-765. | 1-15, 119-134, 274-286, 386-390, 666-703, 749-765. | 1-34, 119-134, 273-286, 388-393, 665-703, 733-765. | 1-41, 118-135, 275-284, 388-397, 664-705, 734-765. |
| TRPC5:EA<sub>4:2</sub>-S2 | 1-15, 119-133, 275-284, 666-700, 750-765. | 1-15, 119-133, 275-284, 629-634, 666-700, 747-765. | 1-15, 119-133, 275-284, 628-632, 666-700, 750-765. | 1-15, 119-133, 275-284, 666-700, 750-765. |

## Supporting Notes

### Supporting Note 1. Determination of TRPC5:EA^PA^ and TRPC5^PA^ structures

Our initial TRPC5:EA^PA^ map (2.7 Å; C4 symmetry) displayed poorly resolved cytosolic domains, indicating structural flexibility. Subsequent 3D variability analysis (3DVA)^49^ in CryoSparc^50^ using ‘clustering mode’ resulted in two well-defined clusters depicting two TRPC5 states (**Figure 1a-f; Extended Data Figure 1a,b,d-g; Extended Data Figure 2**). In state 1 (TRPC5:EA^PA^-S1), the ankyrin repeat domains (ARDs) are close to, and interact with, the coiled-coil domains (CCDs), resembling previous structures of TRPC5 in detergent (PDB 7E4T)^40^ and lipid nanodics (PDB 8GVW)^42^ (**Extended Data Figure 1d-g**). In state 2 (TRPC5:EA^PA^-S2), the ARDs rotate counterclockwise (bottom view) while extending from the symmetry axis, similar to a state of human TRPC5 found in lipid nanodiscs (PDB 7X6C)^42^ (**Extended Data Figure 1d-g**). The final 3D reconstructions, applying C4 symmetry, yielded maps of TRPC5:EA^PA^-S1 and TRPC5:EA^PA^-S2 at global resolutions of 2.5 Å (**Figure 1; Extended Data Figure 2**). The final map of ‘apo’ TRPC5^PA^ (depicting ARD state 2) was obtained at 2.4 Å resolution (C4 symmetry) (**Figure 1; Extended Data Figure 1; Extended Data Figure 2**). Further efforts to classify and sort particles from the ‘apo’ TRPC5^PA^ dataset did not reveal additional states.

### Supporting Note 2. Determination of six distinct TRPC5:EA structures from one data set

To further test the effects of the addition of PA to our samples, including on local maps and on channel states, we decided to try an alternative sample preparation method. Instead of adding an EA/PA preparation to TRPC5 after purification, we maintained EA (100 µM) in all solutions throughout the sample preparation process, from cell lysis to grid making. This resulted in a 2.8 Å cryo-EM map (C4 symmetry; **Extended Data Figure 8**). We observed an unusual density that did not correspond to either EA or the resident lipid, suggesting partial occupancy (**Extended Data Figure 9**). Refining the map without applying symmetry (C1) resulted in a slightly lower resolution map (3 Å) displaying similar density in the EA binding sites, ruling out an artifact arising from symmetrisation.

We next performed 3DVA and visualised the output using ‘simple mode’ in CryoSPARC with 20 frames (**Extended Data Movie 1**). We noted a high degree of variability in the ARDs, consistent with the presence of the two states described for TRPC5:EA^PA^ (see above). Intriguingly, during examination of the area around the EA binding site, we observed a break in symmetry around the pore gate, shifting between C2 and C4 symmetry across different frames and structures (**Extended Data Movie 1**). Because the resolution was capped at 4 Å, this analysis did not provide further detail on EA/lipid stoichiometries in the individual structures. Therefore, we used ‘clustering mode’ to separate the data into the two previously described ARD states (see above). After refinement, we achieved maps with a resolution of ∼3 Å (**Extended Data Figure 8; Extended Data Figure 9**). Although we refined these maps without imposing symmetry (C1), densities in the EA binding sites remained ambiguous.

Further processing with 3DVA allowed us to classify the data from each ARD state into three distinct classes (i.e. resulting in 6 unique maps), which were further categorised based on TRPC5:EA binding stoichiometry (**Extended Data Figure 8; Extended Data Figure 9; Extended Data Figure 11a-f**).

- State 1, full EA occupancy (TRPC5:EA**_4:4_**-S1; 2.8 Å; C4)
- State 1, two EA molecules per TRPC5 tetramer (TRPC5:EA**_4:2-_**S1; 3.0 Å; C4 relaxed symmetry)
- State 1, mixed occupancy, 1-3 EA molecules per TRPC5 tetramer (TRPC5:EA**_mix_**-S1; 3.2 Å; C4 relaxed symmetry)
- State 2, two EA molecules per TRPC5 tetramer (TRPC5:EA**_4:2_**-S2; 3.0 Å; C4 relaxed symmetry)
- State 2, mixed occupancy of 1-3 EA molecules per TRPC5 tetramer (TRPC5:EA**_mix1_**S2; 3.1 Å; C4 relaxed symmetry)
- State 2, mixed occupancy of 1-3 EA molecules per TRPC5 tetramer (TRPC5:EA**_mix2_**-S2; 3.2 Å; C4 relaxed symmetry)

Additional efforts to separate the data for maps with mixed TRPC5:EA stoichiometry were unsuccessful.

## Supplementary Figures

**Supplementary Figure 1.**
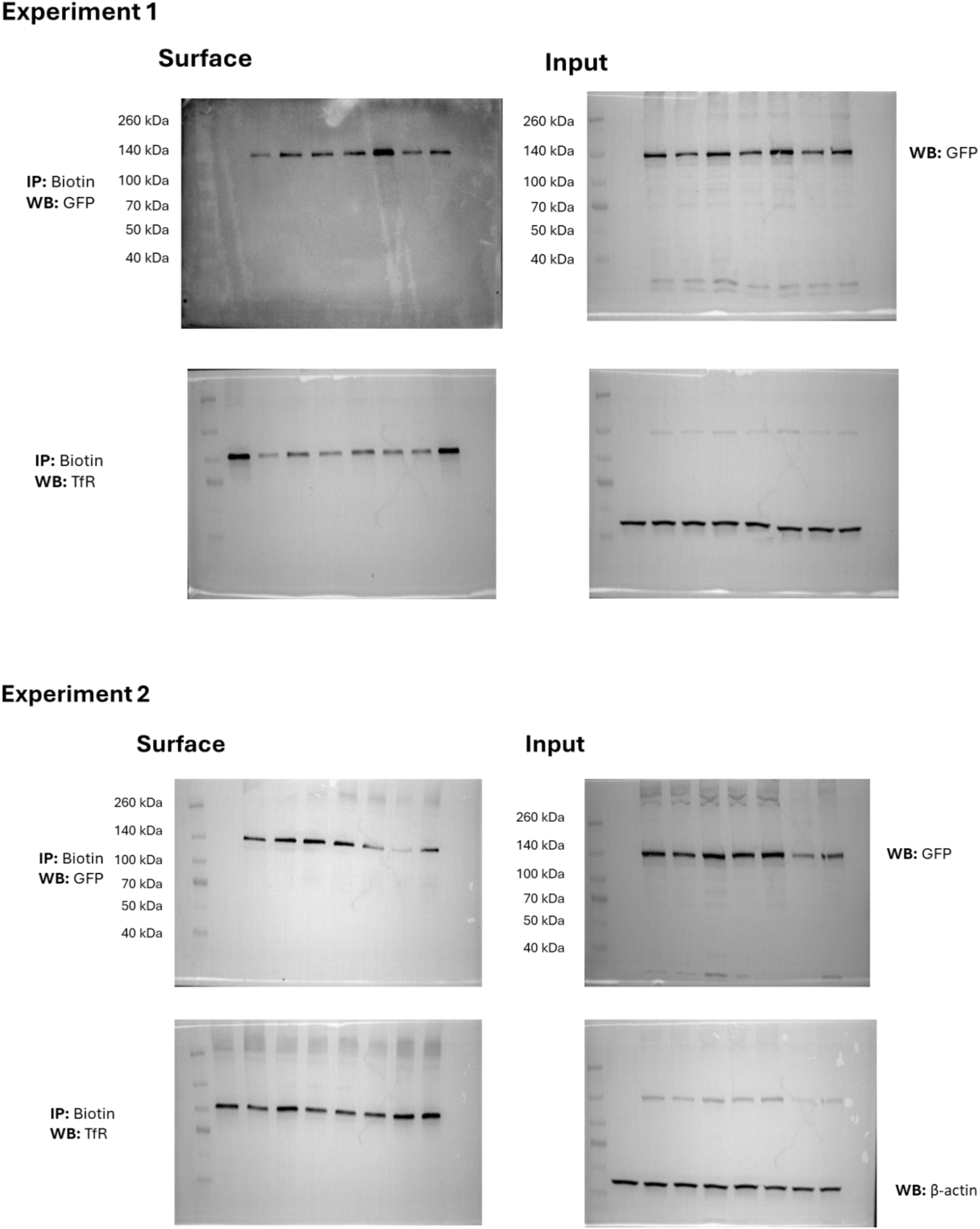
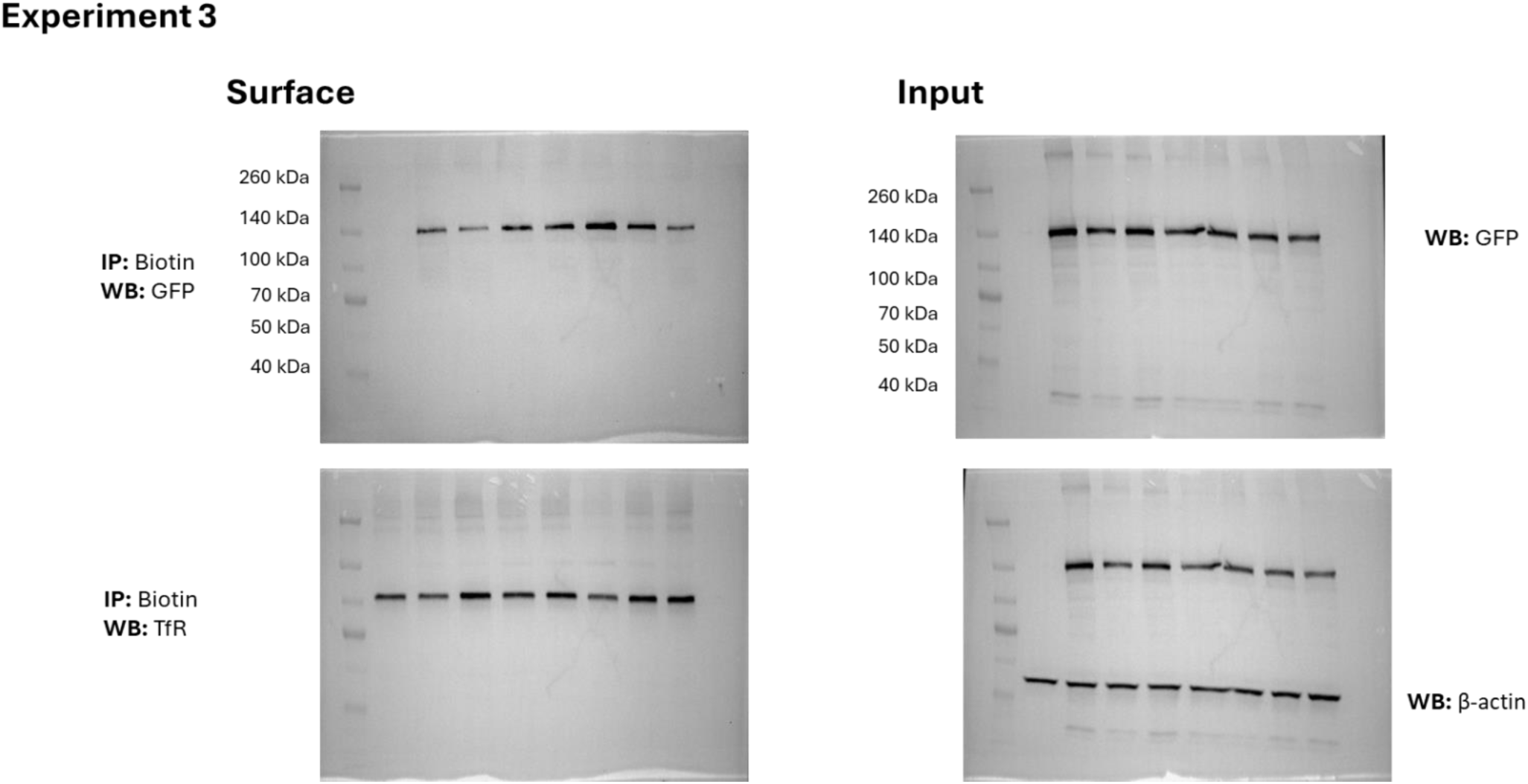
Complete western blots for surface biotinylation experiments described and analysed in Extended Data Figure 5d (three independent experiments).

